# Structural basis of eukaryotic transcription termination by the Rat1 exonuclease complex

**DOI:** 10.1101/2024.03.28.587100

**Authors:** Tatsuo Yanagisawa, Yuko Murayama, Haruhiko Ehara, Mie Goto, Mari Aoki, Shun-ichi Sekine

## Abstract

The 5’-3’ exoribonuclease Rat1 (or Xrn2) plays a pivotal role in termination of mRNA transcription by eukaryotic RNA polymerase II (RNAPII). Rat1 forms a complex with its partner proteins, Rai1 and Rtt103, and acts as a “torpedo” to bind transcribing RNAPII and dissociate DNA/RNA from it. Here we report the cryo-electron microscopy structures of a Rat1-Rai1-Rtt103 complex and three Rat1-Rai1-associated RNAPII complexes (type-1, type-1b, and type-2) from the yeast *Komagataella phaffii*. The Rat1-Rai1-Rtt103 complex structure revealed that Rat1 and Rai1 form a heterotetramer, with a single Rtt103 bound between two Rai1 molecules. In the type-1 complex, Rat1-Rai1 forms a heterodimer and directly binds to the RNA exit site of transcribing RNAPII to extract RNA into the Rat1 nuclease active site. This interaction changes the RNA path in favor of termination (the “pre-termination” state). The type-1b and type-2 complexes have no bound DNA/RNA, likely representing the “post-termination” states. These structures shed light on the mechanism of eukaryotic mRNA transcription termination.

## Introduction

DNA-dependent RNA polymerase (RNAP) carries out gene transcription generally through three steps, initiation, elongation, and termination, which are regulated by specific initiation, elongation, and termination factors ^1,2^. During mRNA transcription by eukaryotic RNA polymerase II (RNAPII), the transcribed pre-mRNA undergoes 5’-capping, splicing, and 3’-polyadenylation, and these processes occur co-transcriptionally ^3^. Assembly of the transcription machineries and their association with processing factors are regulated by post-translational modifications, in particular phosphorylation of the heptad repeat sequence within the C-terminal domain (CTD) of the RNAPII Rpb1 subunit ^4–9^.

Following transcription initiation from the promoter, RNAPII travels along the gene body as a transcription elongation complex (EC) to accomplish transcription of the gene ^10,11^. In the final step of transcription, the EC terminates transcription at a termination site on the gene by dissociating DNA and RNA. Precise control of termination is essential to the maintain/regulate the quality of RNA products, transcriptome, and the genome integrity. Despite its importance, the detailed molecular mechanisms of transcription termination remain largely unknown, especially in eukaryotes. Eukaryotes use multiple mechanisms for transcription termination depending on the type of genes: protein-coding mRNA genes, replication-dependent histone genes, and non-coding RNA genes ^12,13^. Termination of mRNA transcription is a complex process, as it is coupled with mRNA processing. Efficient termination of mRNA transcription requires co-transcriptional cleavage and polyadenylation of the pre-mRNA at the polyadenylation site, followed by 5’-3’ degradation of the downstream nascent RNA transcript to dislodge RNAPII from DNA ^12,14–21^.

The yeast Rat1 protein (Xrn2 in humans) is an essential 5’-3’-exoribonuclease responsible for this very last step ^22–25^. Rat1 forms a binary complex with Rai1, a partner protein that stimulates the Rat1 exoribonuclease activity ^26,27^. Rtt103, a CTD-interaction domain protein, reportedly recruits Rat1-Rai1 to the 3’-end of genes to facilitate EC disassembly ^28,29^. Rat1 is recruited to the monophosphorylated RNA 5’-end newly generated by the pre-mRNA cleavage, and is proposed to act as a “torpedo” to terminate transcription; it digests the RNA emerging from EC in the 5’-3’-direction, catches up with the EC, and dissociates DNA/RNA from RNAPII ^15,22,28,30,31^. The EC disassembly may be accompanied by an allosteric change within the RNAPII structure ^13,32,33^. Nevertheless, the molecular mechanisms by which Rat1 interacts with EC and mediates termination have remained unclear.

In the present study, we analyzed the structures of the Rat1-Rai1-Rtt103 complex and the Rat1-Rai1-bound RNAPII complexes by cryo-electron microscopy (cryo-EM). These structures provide insights into Rat1-Rai1-mediated termination of mRNA transcription by RNAPII.

## Results

### Cryo-EM structure of the Rat1-Rai1-Rtt103 complex

We prepared recombinant proteins of Rat1, Rai1, and Rtt103 from *Komagataella phaffii (K. pastoris).* The purified Rat1 efficiently degrades a 5’-monophosphorylated 31-nt RNA (P-31RNA-Cy5) either in the presence or absence of Rai1 (Supplementary Fig. 1). Mutations of two or five acidic residues in the Rat1 active site (Supplementary Fig. 2) (D233A/D235A (Rat1 2A); E203A/E205A/D233A/D235A/D330A (Rat1 5A); E203Q/E205Q/D233N/D235N/D330N (Rat1 2Q3N)) abolished the nuclease activity of Rat1 (Supplementary Fig. 1B, lanes 6-8). Rai1 or its catalytic mutant (E213A/D215A (Rai1 2A), Supplementary Fig. 3) showed no nuclease activity, consistent with the fact that the substrate for Rai1 is a 5’-triphosphorylated RNA or a 5’-capped RNA (Supplementary Fig. 1B, lanes 9 and 10). In the following structural analyses, we used mutant proteins of Rat1 and Rai1 to avoid RNA digestion.

We prepared the Rat1-Rai1-Rtt103 complex by mixing Rat1 (2A), Rai1(2A), and Rtt103, and determined the cryo-EM structure of the complex at 3.5 Å (Fig. 1, Supplementary Figs. 5-8, and Supplementary Table 1). Although the sample contained a 17-nt RNA, its density was not observed. In this complex, two Rat1-Rai1 heterodimers (dimers 1 and 2) form a tetramer ((Rat1-Rai1)_2_) (Fig. 1). Two Rat1 molecules form interface between dimers 1 and 2. The kinked α12 helix (residues 370-407) of Rat1 in dimers 1 and 2 interacts with the α20 helix (residues 812-819) of Rat1 in dimers 2 and 1, respectively, to form a pseudo twofold symmetric arrangement (Fig. 1A,B,C). We identified the N-terminal domain of Rtt103 (Supplementary Fig. 4) asymmetrically bound between the two Rai1 molecules (Fig. 1A,B,D). Three helices of Rtt103 form extensive interactions with one face of Rai1 in dimer 1, while the N-terminal helix of Rtt103 is close to the same face of Rai1 in dimer 2.

**Fig. 1.**
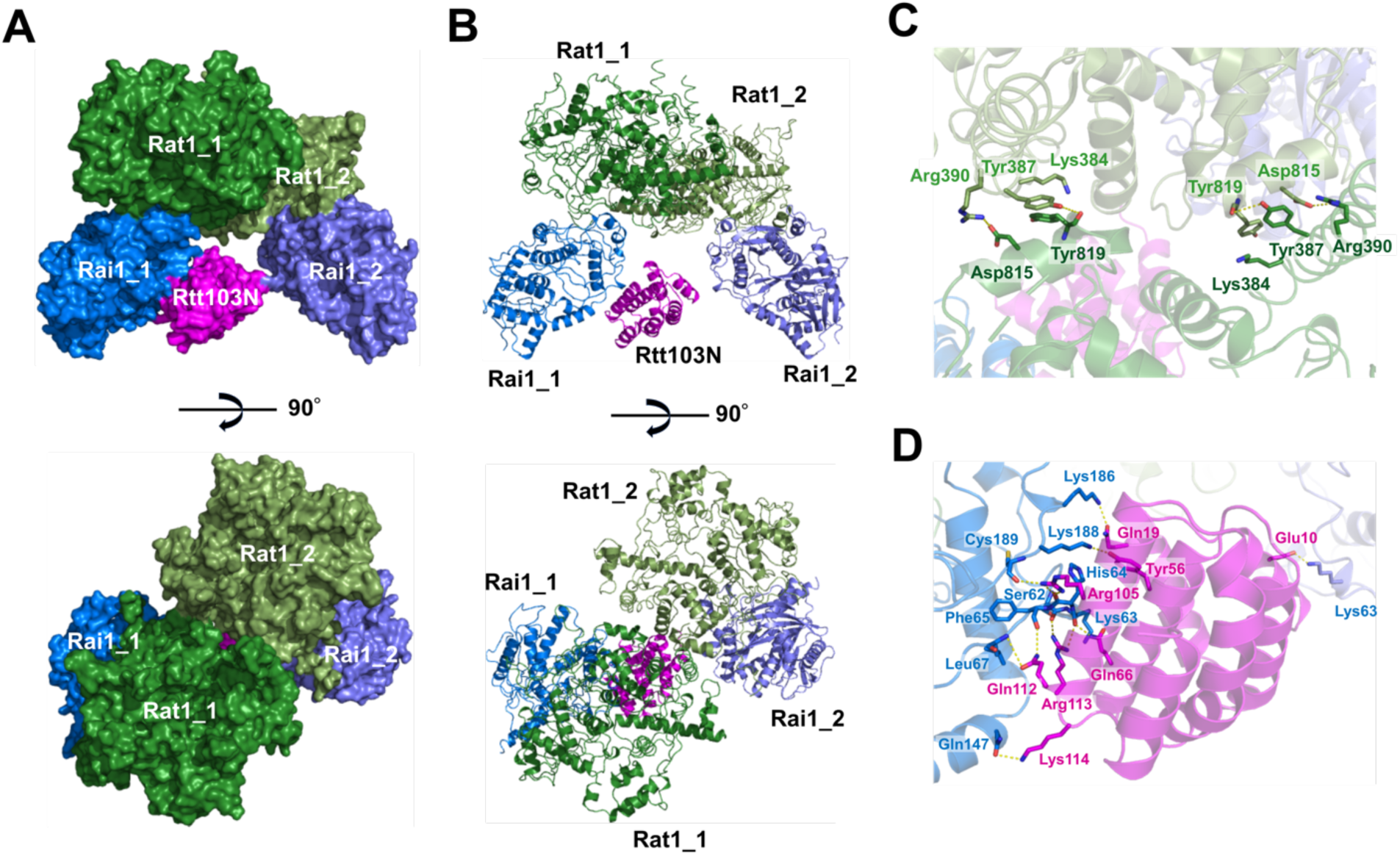
Structure of the Rat1-Rai1-Rtt103 complex. (A, B) The *K. phaffii* Rat1-Rai1 heterotetramer complexed with Rtt103 is represented as a surface and ribbon models. Rat1 is colored green and grass green, Rai1 is colored marine and purple blue, and the N-terminal domain of Rtt103 (Rtt103N) is colored magenta. Bottom, view after a 90° rotation about the horizontal axis from the upper panel. (C) A close-up view of the Rat1 dimer interface. The Rat1 residues involved in dimerization are shown by green stick models. Hydrogen bonding interactions are represented by dotted lines. (D) A close-up view of the interface between Rai1 and Rtt103N. The Rai1 and Rtt103 residues are shown by blue and magenta stick models, respectively. A transparent ribbon model of the Rat1-Rai1 (blue and green) complexed with Rtt103 (magenta) is shown in the background. Hydrogen bonds and ionic interactions between Rai1 and Rtt103 are represented by dotted lines.

### Cryo-EM analysis of the Rat1-Rai1-associated RNAPII complexes

Next, we analyzed the structure of an RNAPII elongation complex (EC) associated with the Rat1-Rai1 exonuclease complex. EC was assembled by mixing *K. phaffii* RNAPII and a DNA/RNA scaffold. Then, to reconstitute the pre-termination complex, the EC was mixed with the Rat1(2Q3N), Rai1(2A), and Rtt103. We examined five RNAs of different length (31 nt, 26, nt, 22 nt, 20 nt, and 17 nt) (Supplementary Fig. 9), and the 22-nt RNA was employed for final data collection, as it yielded the most stable complex. The formed complex species were partially purified by glycerol gradient sedimentation (Supplementary Fig. 10) and crosslinked with BS3 before the preparation of cryogrids. Approximately 21,000 cryo-EM images were collected for single particle analysis (Supplementary Fig. 11, and Supplementary Table 1). Classification of the particles yielded two distinct classes of RNAPII-Rat1-Rai1 complexes (referred to as types 1 and 2), which differ in the Rat1-Rai1 binding mode to RNAPII (Supplementary Figs. 12, 13). In type 1, the Rat1-Rai1 complex is docked around the RNA exit site of the EC. By contrast, in the type-2 complex, the Rat1-Rai1 complex occupies the main channel of RNAPII. The type-1 complex was observed when the RNA length is 20 nt or longer, and the 22-nt RNA yielded the complex in which the Rat1-Rai1 density is best resolved. When the 17-nt RNA was used, particles of the type-1 complex were not observed. These structures are described in detail below.

### Structure of the RNAPII pre-termination complex engaged with Rat1-Rai1 (type 1)

The type-1 complex reveals the Rat1-Rai1 exonuclease complex engaged at the RNA exit site of EC, likely representing the state just prior to transcription termination (Fig. 2). Although Rat1-Rai1 forms a heterotetramer ((Rat1-Rai1)_2_) in the free form, it became a heterodimer (Rat1-Rai1), when bound to RNAPII. Rtt103 was not observed. While Rat1 in the Rat1-Rai1 complex directly interacts with RNAPII, Rai1 is oriented away from RNAPII (Fig. 2A). Rat1 orients its exonuclease active site toward RNAPII to capture the 5’-end of RNA emerging from the RNA exit tunnel of RNAPII. The stalk of RNAPII forms a major interface with Rat1 (Fig. 2B). One side of Rat1 extensively interacts with the Rpb7 subunit of the stalk. While the Rat1 N-terminal loop portion (residues 28-39) fits well to the hydrophobic face of the Rpb7 β-barrel, the C-terminal segment containing Arg715 and Asp 780 contacts the most distal part of the stalk. The segment connecting α5 and β4 (residues 184-186) is also involved in this interaction. Probably because of this interaction, the orientation of the stalk is different as compared with that in EC (Supplementary Fig. 14) ^34^. The Rat1-binding site overlaps with the binding sites of the transcription elongation factors Spt6 and Spt5 (the KOW3-4 domains) ^34,35^, suggesting that Rat1 can access RNAPII after these factors or domains are dissociated from RNAPII (Fig. 2C).

**Fig. 2.**
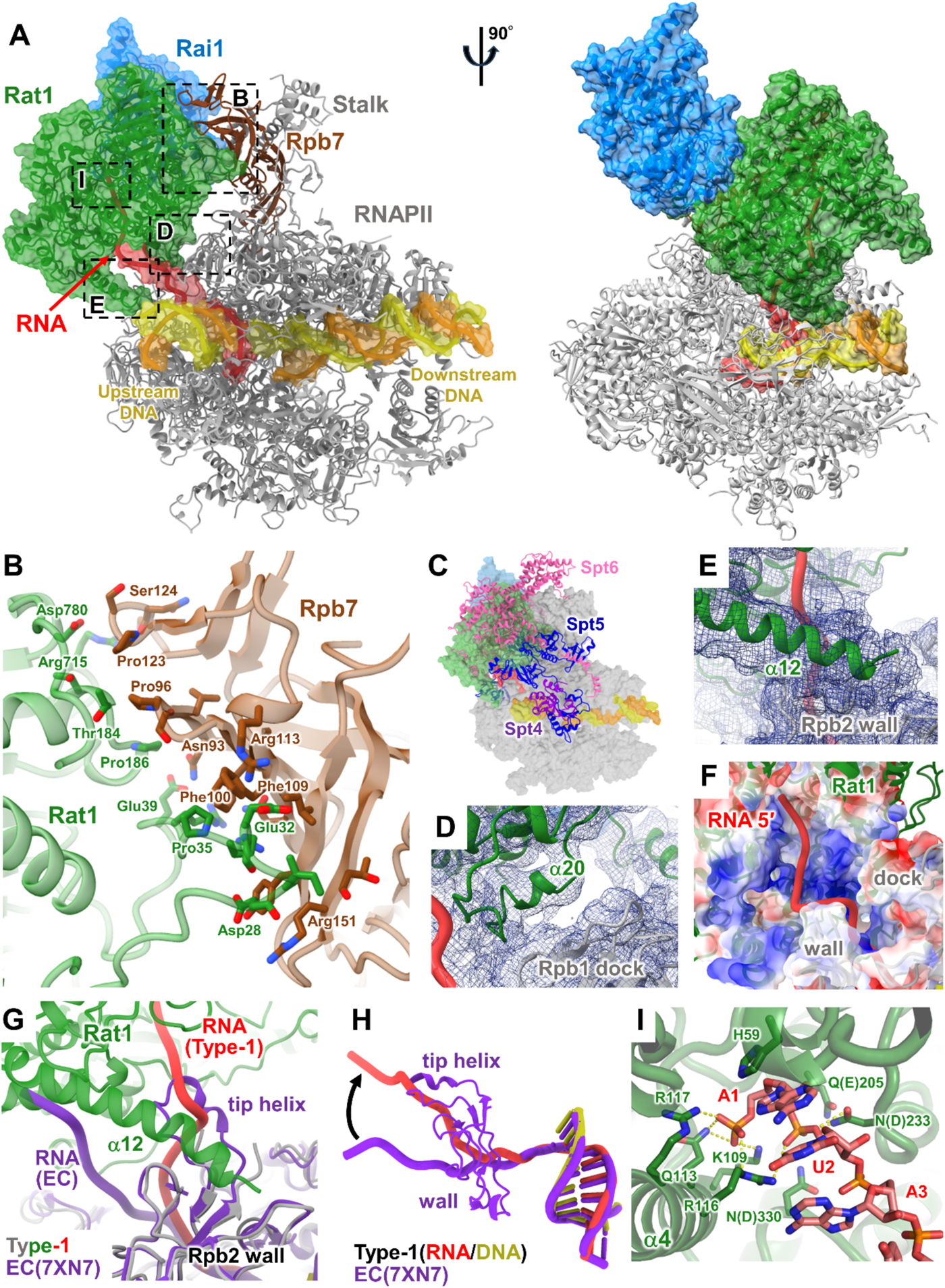
Structure of the RNAPII pre-termination complex (type 1). (A) Structure of EC engaged with Rat1-Rai1 at the RNA exit (the type-1 complex). The structural model is displayed in two orientations. RNAPII, template DNA, non-template DNA, RNA, Rat1, and Rai1 are colored gray (Rpb7 in brown), yellow, orange, green, and blue, respectively. Transparent surfaces of DNA/RNA, Rat1, and Rai1 are overlaid. (B) A close-up view of the Rpb7-Rat1 interface. (C) The Rat1-binding site overlaps with the binding sites of the elongation factors Spt5 and Spt6. The structural model of the type-1 complex is shown in transparent surface, colored as in Fig. 2A. The structural model of RNAPII EC (PDB 7XN7) is superimposed with the type-1 complex by the Rpb1 subunit. Elongation factors Spt4, Spt5 and Spt6 are shown in cartoon representation, and colored purple, blue, and pink, respectively. (D) A close-up view of the interaction between the Rpb1 dock domain and the Rat1 α20 helix. Cryo-EM map (blue mesh) is overlaid on the structural model. (E) A close-up view of the interaction between the Rpb2 wall domain and the Rat1 α12 helix. Cryo-EM map (blue mesh) is overlaid on the structural model. (F) RNA channel in the type-1 complex. A transparent surface of RNAPII and Rat1 (colored by electrostatic surface potentials) is overlaid on the cartoon model (colored as in Fig. 2A). (G, H) Comparison of the RNA path at the RNAPII RNA exit. The structure of RNAPII EC (PDB 7XN7, purple) is superposed on the type-1 complex (colored as in Fig. 2A) by the Rpb1 subunit. (I) Structure of the Rat1 active site with the RNA 5’-end incorporated. Putative hydrogen bonds between Rat1 and RNA are shown as yellow dashed lines. For the residues mutated in the Rat1 active site (residues 205, 233, and 330), the wild-type amino acids are shown in parentheses.

In addition to the stalk interaction, the α20 helix (residues 812-819) of Rat1 contacts the dock domain of the Rpb1 subunit of RNAPII (Fig. 2D). The kinked α12 helix (residues 385-407) also contacts the wall domain ^36^ of the Rpb2 subunit of RNAPII (Fig. 2E). Probably because of this interaction, the tip helix of the wall domain is displaced and disordered. These interactions form a continuous basic channel connecting the RNAPII RNA exit with the Rat1 active site (Fig. 2F). The RNA segment emerging from the RNA exit tunnel assumes an extended conformation, and is accommodated in the channel. Consequently, the RNA path is largely different from that in EC (Fig. 2G, H). The 5’-terminal three nucleobases (A1-U2-A3) are stacked to each other, and the 5’-terminal nucleobase (A1) is stacked with His59 at the bottom of the active site (Fig. 2I, Supplementary Fig. 15). The 5’-phosphate group of A1 is specifically recognized by Lys109, Gln113, Arg116, and Arg117 on the tower helix (α4) at the bottom of the active site ^37^. The scissile phosphate bond also contacts Lys109 and Arg116, and is close to the putative catalytic residues Glu205, Asp233, and Asp330, which are mutated to Gln or Asn to avoid RNA cleavage. Thus, Rat1 is docked to the RNA exit of RNAPII to extract RNA into its active site, largely changing the RNA path.

Particle classification identified another structural class similar to the type-1 complex, which is referred to as type-1b (Fig. 3). In this complex, the RNAPII conformation and the position of the bound Rat1-Rai1 complex are similar to those in the type-1 complex. However, the type-1b complex lacks clear nucleic-acids densities within RNAPII, leaving the RNAPII main channel and RNA-exit channel empty. Despite the lack of DNA/RNA, the RNAPII clamp is in a closed conformation as in EC. We presume that the type-1b complex was formed after the release of DNA/RNA from the type-1 complex via a transient conformational change of RNAPII such as opening of the clamp, due to the binding of Rat1-Rai1.

**Fig. 3.**
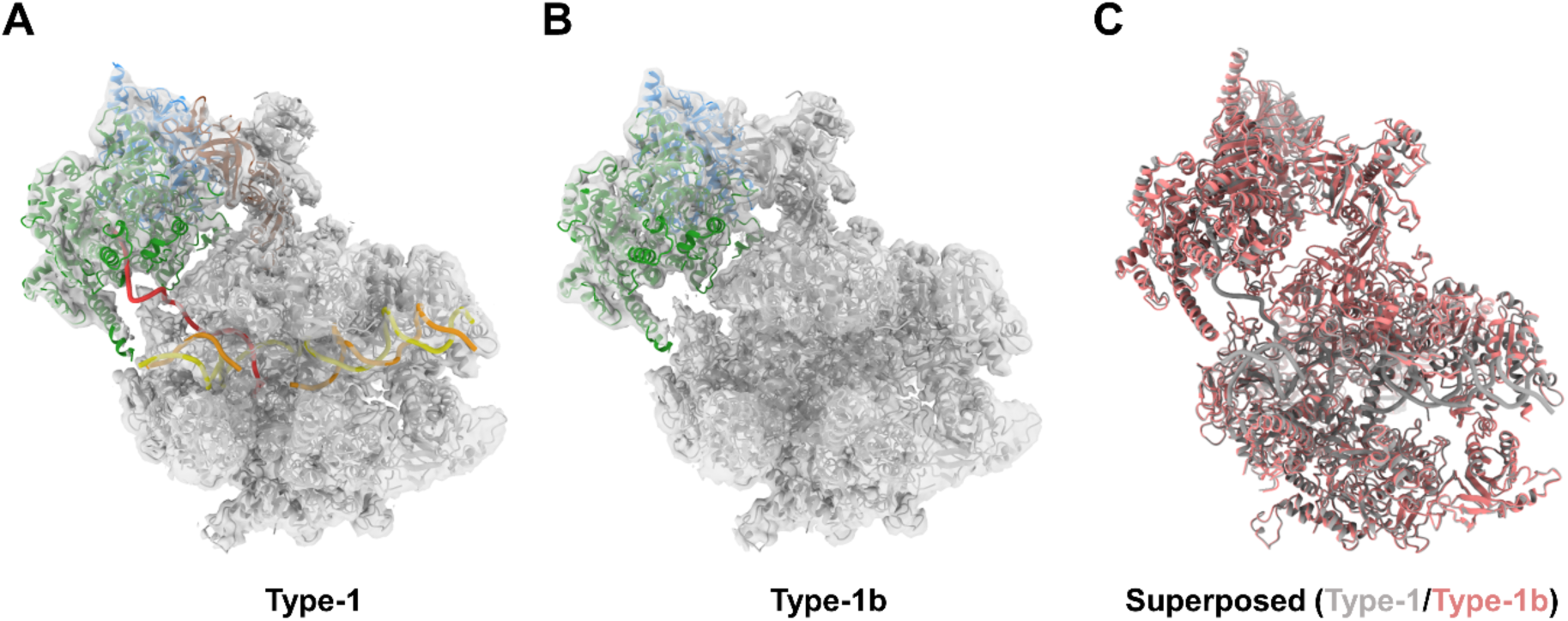
Structure of RNAPII bound with Rat1-Rai1 at the RNA exit (type 1b). (A, B) Structures of the type-1 and type-1b complexes. Cryo-EM maps (white surfaces) are overlaid on the structural models (cartoon representation, colored as in Fig. 2). (C) Comparison of the type-1 and type-1b complexes. The structural models for the type-1 and type-1b complexes are superimposed by the Rpb1 subunit.

### Structure of the Rat1-Rai1 complex bound within the RNAPII cleft (type 2)

We also identified a different structural class (type 2), in which the Rat1-Rai1 complex is bound to RNAPII in a different manner than the type-1 complex. In the type-2 complex, the Rat1-Rai1 complex occupies the main cleft of RNAPII, instead of DNA/RNA (Fig. 4A-C). While Rat1 is sandwiched between the Rpb1 clamp head domain and the Rpb2 lobe domain, Rai1 is sandwiched between the Rpb1 clamp domain and the Rpb2 protrusion domain (Fig. 4D,E). Rat1 is also close to the Rpb5 subunit of RNAPII. The Rat1 α20 helix interacts with the basic face of the Rpb2 lobe (Fig. 4D). The Rat1-Rai1 binding site overlaps with the binding site of the double-stranded DNA in the transcription pre-initiation complex (PIC) ^38,39^ (Fig. 4F) and the binding sites of the elongation factors Elf1, Spt5 (the NGN and KOW1 domains), and Spt4 in EC ^40,41^ (Fig. 4G). Rat1-Rai1 may preferentially bind to the RNAPII cleft, when RNAPII releases DNA/RNA from the cleft. The type-2 Rat1-Rai1 binding may prevent non-specific DNA binding by RNAPII, until it is recycled into a PIC to initiate another round of transcription.

**Fig. 4.**
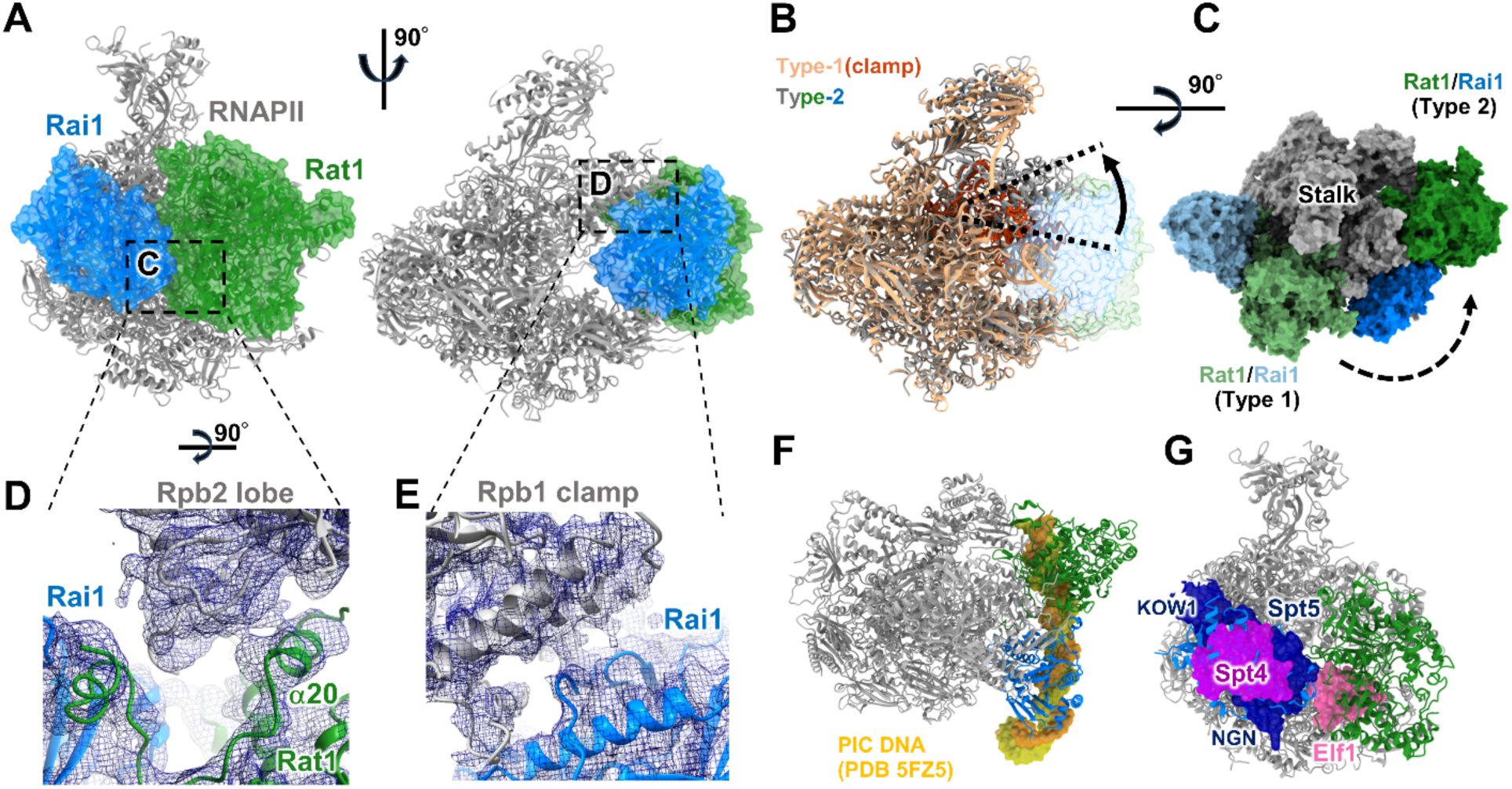
Structure of Rat1-Rai1 bound within the RNAPII cleft (type 2). (A) Structure of Rat1-Rai1 bound within the RNAPII cleft (the type-2 complex). The structural model is displayed in two orientations, colored as in Fig. 2. Transparent surfaces of Rat1 and Rai1 are overlaid. (B) Comparison of the type-1 and type-2 complexes. The structural models for the type-1 (light orange) and type-2 (colored as in Fig. 4A) complexes are superimposed by the Rpb2 subunit. Structural models for the RNAPII are shown in cartoon representation. The RNAPII clamp of the type-1 complex is colored red. Structural models for Rat1-Rai1 in the type-2 complex is shown as a transparent surface. (C) Comparison of the Rat1-Rai-1 binding sites in the type-1 and type-2 complexes. The structural models are superimposed by the Rpb2 subunit. The structural models for the type-2 complex (colored as in Fig. 4A) and Rat1-Rai1 of the type-1 complex (colored light green and light blue) are shown in surface representation. (D) A close-up view of the interaction between the Rpb2 lobe and the Rat1 α20 helix. Cryo-EM map (blue mesh) is overlaid on the structural model. (E) A close-up view of the interaction between the RNAPII clamp and the Rai1. Cryo-EM map (blue mesh) is overlaid on the structural model. (F) Overlapping binding sites for Rat1-Rai1 and double-stranded DNA in the transcription pre-initiation complex (PIC). The structural model of the type-2 complex is shown in cartoon representation, colored as in Fig. 4A. The structural model of RNAPII PIC (PDB 5FZ5) is superimposed on the type-2 complex by the Rpb2 subunit. The template DNA and the non-template DNA of the PIC are shown in surface representation, and colored yellow and orange, respectively. (G) Overlapping binding sites for Rat1-Rai1 and elongation factors Spt4, Spt5, and Elf1. The structural model of the type-2 complex is shown in cartoon representation, colored as in Fig. 4A. The structural model of RNAPII EC (PDB 7XN7) is superposed on the type-2 complex by the Rpb2 subunit. Elongation factors Spt4, Spt5, and Elf1 are shown in surface representation, and colored purple, blue, and pink, respectively.

## Discussion

In this study, we determined the cryo-EM structure of the termination factor complex Rat1-Rai1-Rtt103 from the yeast *K. phaffii*. Two Rat1-Rai1 dimers form a tetramer ((Rat1-Rai1)_2_) mediated by the Rat1-Rat1 interaction, and Rtt103 is bound between two Rai1 molecules. We also determined the structures of RNAPII associated with the Rat1-Rai1 heterodimer complex. Rat1-Rai1 revealed two alternative RNAPII binding modes (types 1 and 2). In the type-1 complex, Rat1-Rai1 binds to the RNAPII stalk, orienting the exonuclease active site of Rat1 toward the RNA-exit site of EC. Rat1 forms direct contacts with the RNAPII dock and wall domains, and incorporates the RNA 5’-end emerging from the RNA exit. This Rat1 interaction changes the direction of RNA, as compared with that in EC. As the type-1 complex was observed with RNAs longer than 20 nt, this state likely represents the “pre-termination” state. Consistently, we also identified the type-1b complex, in which DNA/RNA is dissociated from RNAPII (the “post-termination” state). On the other hand, in the type-2 complex, Rat1-Rai1 is bound within the widely-opened RNAPII main cleft. This structure could represent another post-termination state. These structures outline the mechanism of eukaryotic mRNA transcription termination (Fig. 5). (i) Rat1-Rai1 exists as a tetramer when it is unbound to RNAPII, but the tetramer dissociates into a dimer when bound to RNAPII. When cleavage of mRNA precursor occurs at the polyadenylation site, Rat1 captures the mono-phosphorylated RNA 5’-end, and starts trimming of the RNA to track RNAPII. (ii) When Rat1 trimmed the RNA to its length of 20∼22 nt, Rat1-Rai1 forms a close contact with RNAPII. The Rat1 binding to EC requires prior dissociation of Spt6 and Spt5 (KOW3-4) from the EC. Further RNA trimming would increase tension of RNA between the Rai1 and RNAPII active sites, leading to perturbation/destabilization of the DNA/RNA hybrid within the transcription bubble or in the RNAPII-DNA/RNA contacts. (iii) This would trigger bubble collapse and release of DNA/RNA. (iv) After the termination, Rat1-Rai1 is relocated to the empty DNA-binding site of RNAPII to occlude it. Alternatively, Rat1-Rai1 could move from the RNA-exit site to the RNAPII cleft, while digesting RNA to actively dislodge the DNA/RNA from the RNAPII cleft. The Rat1-Rai1 bound to the RNAPII cleft would protect RNAPII, until the RNAPII is recycled into a PIC for another round of transcription.

**Fig. 5.**
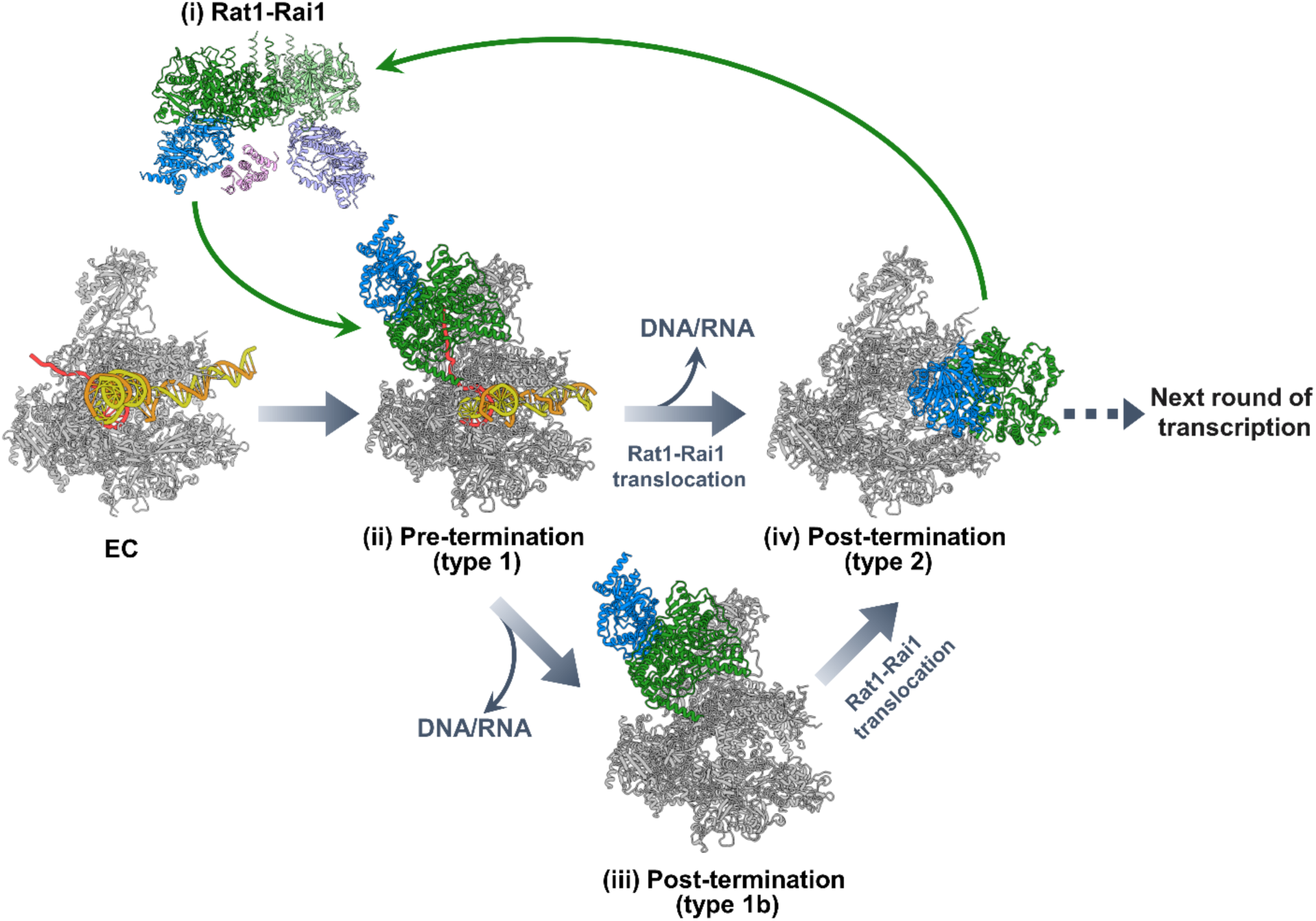
Model of Rat1-dependent transcription termination.

Rat1 is reportedly involved not only in mRNA transcription by RNAPII, but also in rRNA transcription by RNA polymerase I (RNAPI) ^42,43^. Superimposition of the current type-1 complex structure with the RNAPI EC structure showed no apparent steric clash between Rat1 and RNAPI, except for the RNAPI stalk (Supplementary Fig. 16). If the orientation of the RNAPI stalk changes like that of RNAPII, Rat1 could bind RNAPI as in the same way as the type-1 complex. In contrast, the stalk of RNA polymerase III (RNAPIII) has specific insertions, which could make Rat1 binding incompatible.

Rat1 binds at the RNA-exit site of RNAPII to incorporate RNA into its active site. This view is analogous to the manner in which the bacterial termination factor Rho interacts with transcribing RNAP ^44,45^. Although Rho is an ATP-dependent RNA helicase/translocase, it binds the similar part of RNAP (the β-flap and zinc-binding domains) to extract nascent RNA. Although these enzymes differ in their molecular structures, enzymatic activities, and detailed RNAP interactions, they may share a similarity in that they are dedicated to bind both the nascent RNA and RNAP to destabilize EC, suggesting a mechanistic generality of transcription termination.

The Rat1/Xrn2 action in mRNA transcription termination is tightly coupled with prior cleavage of the mRNA precursor and polyadenylation by the cleavage and polyadenylation (CPA) complex ^12^. Further studies are needed to clarify how Rat1/Xrn2 cooperates with the CPA complex at appropriate timing to complete transcription termination coupled with mRNA 3’-processing.

## Methods

### Materials

Biochemical and molecular biological procedures were performed with commercially available materials, enzymes, and chemicals. Synthesized DNAs and RNAs were purchased from Eurofins Genomics (Tokyo, Japan).

### Cloning, expression, and purification of *kp*Rat1 proteins

The DNA fragment encoding *Komagataella phaffii* Rat1 (*kp*Rat1) gene was PCR-amplified from a plasmid containing a codon-optimized *kp*Rat1 gene as a template. The *kp*Rat1 gene, and a maltose-binding protein (MBP) gene with a human rhinovirus 3C (HRV-3C) cleavage site at the N-terminus, and a hexahistidine (6×His)-tag at the C-terminus were cloned into the pET47 vector. The *E. coli* Rosetta2 (DE3) cells (Novagen) were transformed with the plasmid and selected on an LB agar plate supplemented with 50 μg/ml kanamycin. Single colonies were used to inoculate 10 mL of Plusgrow II culture media (Nacalai), supplemented with 50 μg/ml kanamycin, and grown at 37°C for 4 hrs. The starter culture was transferred to two-liter flasks of the Plusgrow II culture, and then grown at 37°C. When the OD_600_ reached 0.6, the temperature was lowered to 20°C. The protein expression was induced with 1 mM IPTG, and the cells were grown at 20°C for 21 hrs. The *E. coli* cells were collected by centrifugation and stored at –80°C. The cells were resuspended in 50 mM potassium phosphate buffer (pH 7.4), containing 500 mM NaCl, 25 mM imidazole, 5 mM β-mercaptoethanol, and protease inhibitor cocktail (Complete-EDTA free ULTRA, Roche) (buffer A), and were lysed by sonication on ice. The cell lysate was centrifuged at 15,000 × g for 15 min at 4 °C, and the supernatant was applied to a HisTrap column (Cytiva), which was equilibrated with buffer A. The *kp*Rat1-MBP-6×His fusion protein was eluted with buffer A containing 400 mM imidazole, instead of 25 mM imidazole, and collected. The eluted fraction was collected and dialyzed against 40 mM potassium phosphate buffer (pH 7.4), containing 50 mM NaCl and 1 mM DTT (buffer B). The dialyzed fraction was then applied to a Resource Q column (Cytiva), and after washing the column with buffer B, the bound proteins were eluted by a linear gradient of 50–537.5 mM NaCl. The C-terminal MBP-6xHis-tag was cleaved with HRV-3C protease at 4 °C overnight. The dialyzed fraction was then loaded on a HiTrap Heparin column (Cytiva), and after washing the column with buffer B, the bound *kp*Rat1 proteins were eluted by a linear gradient of 50–1025 mM NaCl. The eluted fractions were collected, dialyzed against 20 mM HEPES-NaOH buffer (pH 7.5), containing 200 mM NaCl, and 10 mM β-mercaptoethanol (buffer C), and concentrated by ultracentrifugation. Aliquots of the *kp*Rat1 protein were flash-cooled in liquid nitrogen and stored at –80 °C. The catalytically inactive mutant *kp*Rat1 proteins, *kp*Rat1(E203Q/E205Q/D233N/D235N/D330N)[*kp*Rat1(2Q3N)], *kp*Rat1(E203A/E205A/D233A/D235A/D330A)[*kp*Rat1(5A)], and *kp*Rat1(D213A/D215A)[*kp*Rat1(2A)], were expressed and purified in the same manner as the wild-type *kp*Rat1 protein.

### Cloning, expression, and purification of *kp*Rai1 proteins

The DNA fragment encoding *K. phaffii* Rai1 (*kp*Rai1) gene was PCR-amplified from *K. phaffii* genomic DNA and cloned into the modified pCR2.1 vector possessing a 6×His tag and an HRV-3C cleavage site attached at its N-terminus. The *E. coli* KRX cells (Promega) were transformed with the plasmid and selected on an LB agar plate supplemented with 50 μg/ml kanamycin. Single colonies were inoculated into 10 mL of Plusgrow II culture media (Nacalai), supplemented with 50 μg/ml kanamycin, and grown at 37°C for 4 hrs. The starter culture was transferred to one-liter flasks of the Plusgrow II culture, and then grown at 37°C. When the OD_600_ reached 0.5, the temperature was then lowered to 20°C. The protein expression was induced with 0.2% rhamnose, and the cells were grown at 20°C for 21 hrs. The *E. coli* cells were collected by centrifugation and stored at –80°C. The cells were resuspended in buffer A, and were lysed by sonication on ice. The cell lysate was centrifuged at 15,000 × g for 15 min at 4°C, and the supernatant was applied to a HisTrap column (Cytiva), which was equilibrated with buffer C. The protein was eluted with buffer A containing 400 mM imidazole, instead of 25 mM imidazole, and collected. The eluted fraction was collected and dialyzed against buffer B. The dialyzed fraction was then applied to a Resource Q column (Cytiva), and after washing the column with buffer B, the bound proteins were eluted by a linear gradient of 50–537.5 mM NaCl. The N-terminal 6×His-tag was cleaved with HRV-3C protease at 4 °C overnight. The dialyzed fraction was then loaded on a HiTrap Heparin column (Cytiva), and after washing the column with buffer B, the bound proteins were eluted by a linear gradient of 50–1025 mM NaCl. The eluted fractions were collected, dialyzed against buffer C, and concentrated by ultracentrifugation. Aliquots of the purified *kp*Rai1 protein were flash-cooled in liquid nitrogen and stored at –80 °C. The catalytically-inactive mutant *kp*Rai1 protein, *kp*Rai1(E213A/D215A)[*kp*Rai1(2A)], was expressed and purified in the same manner as the wild-type *kp*Rai1 protein.

### Cloning, expression, and purification of *kp*Rtt103 protein

The DNA fragment encoding *K. phaffii* Rtt103 (*kp*Rtt103) gene was PCR-amplified from *K. phaffii* genomic DNA and cloned into the modified pCR2.1 vector with an HRV-3C cleavage site and a 6×His tag attached at its C-terminus. The *E. coli* KRX cells were transformed with the plasmid and selected on an LB agar plate supplemented with 50 μg/ml kanamycin. Single colonies were inoculated into 10 mL of Plusgrow II culture media (Nacalai), supplemented with 50 μg/ml kanamycin, and grown at 37°C for 4 hrs. The starter culture was transferred to one-liter flasks of the Plusgrow II culture, and then grown at 37°C. When the OD600 reached 0.6, the temperature was then lowered to 20°C. The protein expression was induced with 0.2% rhamnose, and the cells were grown at 20°C for overnight. The *E. coli* cells were collected by centrifugation and stored at –80°C. The cells were resuspended in buffer A, and were lysed by sonication on ice. The cell lysate was centrifuged at 15,000 × g for 15 min at 4 °C, and the supernatant was applied to a HisTrap column (Cytiva), which was equilibrated with buffer A. The protein was eluted with buffer A containing 400 mM imidazole, instead of 25 mM imidazole, and collected. The eluted fraction was collected and dialyzed against buffer B. The dialyzed fraction was then applied to a Resource Q column (Cytiva), and after washing the column with buffer B, the bound proteins were eluted by a linear gradient of 50–537.5 mM NaCl. The N-terminal 6×His-tag was cleaved with HRV-3C protease at 4 °C overnight. The dialyzed fraction was then loaded on a HiTrap Heparin column (Cytiva), and after washing the column with buffer B, the bound proteins were eluted by a linear gradient of 50–1025 mM NaCl. The eluted fractions were pooled, concentrated, and applied to a HiLoad 16/60 Superdex 200 column (Cytiva), equilibrated with 30 mM potassium phosphate buffer (pH 7.4), containing 200 mM NaCl and 1 mM DTT. The eluted fractions were collected, dialyzed against buffer C, and concentrated by ultracentrifugation. Aliquots of the purified *kp*Rtt103 protein were flash-cooled in liquid nitrogen and stored at –80 °C.

### Purification of RNA polymerase II from *K. phaffii*

*kp*RNAPII containing the TAP-tagged Rpb2 subunit was purified from a genetically modified strain of *K. phaffii*, as described previously ^46,47^, with the exception that the Ni-column chromatography step was omitted and the sample was dialyzed against RNAPII buffer [20 mM Tris-acetate (pH 8.0), 150 mM potassium acetate, 1 μM zinc acetate, 0.1 mM tris(2-carboxyethyl)phosphine (TCEP) and 5% glycerol] before concentration.

### *In vitro* assay of the 5’ → 3’-exoribonuclease activity

*kp*Rat1/Rai1 activity assays were carried out in 10-μl reaction mixtures containing 20 mM HEPES-NaOH (pH 7.5), 100 mM NaCl, 5 mM MgCl_2_, 5 mM DTT, and 0.75 μM 5’-phosphorylated/3’-Cy5-labelled 31 base RNA substrate (5’-P-UAAUCCCAUAUAUAUGCAUAAAGACCAGGCU-Cy5-3’)(P-31RNA-Cy5). The purified *kp*Rat1 (1.5 μM) and *kp*Rai1 (1.5 μM) proteins were mixed, and incubated at 0°C for 1hr. The reaction was terminated by the addition of 1 volumes of 2 x SDS loading buffer (0.125 M Tris-HCl (pH 6.8), 4% SDS, 10% glycerol, 10% β-mercaptoethanol, 0.03% Orange G) and heat treatment (95°C for 3min). Reaction products were resolved on 10-20% SDS-PAGE gels and visualized using LAS4000 (Leica), and stained with Simply Blue (Thermo Fisher Scientific).

### Preparation of the Rat1-Rai1-Rtt103-RNA complex for cryo-EM analysis

5’-Cy5-labelled 17 base RNA (5’-Cy5-CTAGTAATGACCAGGCU-3’)(Cy5-17RNA), *kp*Rat1(2A), *kp*Rai1(2A), and *kp*Rtt103 were mixed in a 2:1:2:2 molar ratio (15 μM, 7.5 μM, 15 μM, and 15 μM, respectively) in 20 mM HEPES-NaOH buffer (pH 7.5), containing 100 mM NaCl, 5 mM MgCl_2_, and 2 mM DTT, and incubated on ice for 30 min to allow for association. After filtration, the sample was loaded and purified over a Superose 6 Increase 10/300 GL column (Cytiva) in 20 mM HEPES-NaOH buffer (pH 7.5), containing 100 mM NaCl, 1 mM MgCl_2_, and 1 mM DTT. The peak corresponding to the complex was pooled, concentrated by centrifugal filtration (Amicon), and buffer exchanged into 20 mM HEPES-NaOH buffer (pH 7.5), containing 100 mM NaCl, 1 mM MgCl_2_, and 1 mM TCEP. 5-10 μl of the protein-nucleic acid complexes were crosslinked with 5 mM Bissulfosuccinimidyl suberate (BS3) (Pierce) on ice for 30 min, and quenched with 100 mM Tris-HCl (pH7.5).

### Preparation of RNAPII bound with Rat1-Rai1-Rtt103 for cryo-EM analysis

To assemble EC bound with the Rat1-Rai1-Rtt103 complex for cryo-EM studies, purified *kp*Rat1(2Q3N), *kp*Rai1(2A), and *kp*Rtt103 were incubated with pre-assembled *kp*RNAPII-DNA/RNA-scaffold complex in buffer containing 20 mM HEPES-NaOH (pH7.5), 100 mM NaCl, 10 mM KCl, 1 mM MgCl_2_, and 1 mM DTT. The scaffold in use is a DNA-RNA scaffold, consisting of a non-template DNA (5’-GTCAAGGCAGTACTAGTAATTTAGCAATCCAACTACTTTATCTTTTAATCAATCTACAATAACTGGGGGGCTACCGACGCTAGGGATCCT-3’), a template DNA (5’-AGGATCCCTAGCGTCGGTAGCCCCCCAGTTATTGTAGATTGATTAAAAGATAAAGTAGTTGAGCCTGGTCATTACTAGTACTGCCTTGAC-3’), and a 5′-phosphorylated/3’-Cy5-labelled 22base RNA (5’-P-AUAUAUGCAUAAAGACCAGGCU-Cy5-3’) (P-22RNA-Cy5) (Supplementary Fig. 9). The template and non-template DNAs were designed to have a 9 bp mismatch, so the scaffold contains a stable DNA-RNA hybrid in the transcription-bubble. The P-22RNA-Cy5, template DNA, and non-template DNA were mixed and annealed at 95°C for 2 min in annealing buffer (10mM HEPES-NaOH (pH 7.5), 50 mM NaCl), and gradually decreasing the temperature from 95°C to room temperature in the buffer. *kp*RNAPII (1.4 μM) was mixed with the scaffold (1.6 μM), and incubated on ice for 1 hr in RNAPII complex buffer (20 mM HEPES-NaOH buffer (pH7.5), containing 100 mM NaCl, 10mM KCl, 1 mM MgCl_2_, and 5 mM DTT). Then, *kp*Rat1(2Q3N) (8 μM), *kp*Rai1(2A) (8 μM), and *kp*Rtt103 (8 μM) were added, and incubated on ice for 6 hrs to allow for association.

The protein-nucleic acid complexes were then fractionated by ultracentrifugation sedimentation on a glycerol gradient. The glycerol gradient (20 mM HEPES-NaOH buffer (pH 7.5), containing 100 mM NaCl, 10 mM KCl, 1 mM MgCl_2_, 5 mM DTT, 10-30% glycerol) was prepared by layering from the lowest glycerol concentration solution to the highest glycerol concentration solution from the bottom of the tube with a syringe. The protein-nucleic acid complexes were applied on the top of the gradient solution and ultracentrifuged at 33,000 rpm at 4 °C for 17 hr for 45 min, using an S55S rotor (Eppendorf Himac Technologies). The fractionated samples containing the *kp*RNAPII/DNA-RNA scaffold/Rat1(2Q3N)/Rai1(2A)/Rtt103 complexes were pooled, and concentrated with Amicon Ultra 50K centrifugal filters (Millipore), and buffer exchanged to 20 mM HEPES-NaOH buffer (pH 7.5), containing 100 mM NaCl, 10 mM KCl, 1 mM MgCl_2_, and 1 mM TCEP. Crosslinking of the protein-nucleic acid complexes (5-10 μl) was performed with 5 mM BS3 on ice for 40 min, and quenched with 100 mM Tris-HCl (pH7.5).

### Cryo-EM sample preparation and data collection

2.5-3.0 μL of the crosslinked complexes were applied onto 300-mesh Quantifoil R1.2/1.3 Cu grids (Quantifoil Micro Tools GmbH, Germany), which were glow-discharged before use with a PIB-10 ION Bombarder (Vacuum Device Inc.). The grids were then blotted for 3.0 s at 10°C and 75% humidity and plunge-frozen in liquid ethane using an EMGP2 (Leica, Germany). Cryo-EM data were collected with a 300 kV Krios G4 transmission electron microscope (Thermo Fisher Scientific, USA) equipped with a BioQuantum energy filter and K3 camera (Gatan, USA). Images were recorded at 105,000 × magnification with a calibrated pixel size of 0.83 Å/pixel. The exposure times were set to 2.2-2.4 s with a total accumulated dose of 60-62 electrons per Å^2^. All images were automatically recorded using the EPU software (Thermo Fisher Scientific, USA). For the *kp*Rat1(2A)/Rai1(2A)/Rtt103 complex, a total of 11,000 images were collected with a defocus range from −2.4 μm to −1.6 μm. For the *kp*RNAPII/DNA-RNA scaffold/Rat1(2Q3N)/Rai1(2A) complex, a total of 21,135 images were collected with a defocus range from −2.4 μm to −1.6 μm. Statistics for data collection and refinement are in Supplementary Table 1.

### Image processing and 3D reconstruction of the Rat1-Rai1-Rtt103 complex

The image processing of *kp*Rat1(2A)/Rai1(2A)/Rtt103 data was performed with Relion 3.1, unless otherwise specified ^48,49^. After motion correction, the CTF parameters were estimated with CTFFIND ^50^, and then, the initial particles were picked with Relion blob picker. After preliminary 2D and 3D classifications, a Topaz network was trained with good particles, and then particles were re-picked with Topaz ^51^. Repeated cycles of 2D classification, 3D classification, 3D refinement, Bayesian-polishing and CTF refinement were performed to select good particles and to improve the resolution (Supplementary Fig. 6). C2 symmetry was applied to the most of these steps. Finally, 3.47 Å map of *kp*Rat1/Rai1 was obtained from 526,250 particles, also with C2 symmetry applied. To resolve the Rtt103N density between the dimers, final steps of 3D classifications were done without symmetry, and with local angular searches and the relax symmetry (C2) options. Then, the final 3.78 Å map of *kp*Rat1/Rai1/Rtt103N was obtained from 77,924 particles

### Image processing and 3D reconstruction of the RNAPII-Rat1-Rai1 complex

Motion correction was performed with Relion3.1 ^48,49^, and the CTF parameters were estimated with CTFFIND ^50^. Initial particle picking was performed with Warp ^52^ and Topaz ^51^, and the particle coordinates from each program were merged after 2D classification. Approximately 1,500,000 particles containing RNAPII were selected by 2D and 3D classifications, and subjected to 3D refinement followed by local CTF estimation, beam tilt refinement and Bayesian polishing to improve the resolution. The particles were further sorted by 3D classifications with blush regularization using relion5 ^53^ into apo RNAPII, EC and RNAPII-Rat1-Rai1 complexes (type 1 and type 2) (Supplementary Fig. 12). The 3D classes of EC, which showed weak density of Rat1-Rai1 at the type-1 binding site, were subjected to focused 3D classification with a mask around Rat1-Rai1 and the RNAPII stalk. The particles with clear Rat1-Rai1 density were merged with the type-1 particles from preceding 3D classification. The particle picking network of Topaz was trained on the particle coordinates of type-1 and type-2 complexes, and the particles were re-picked with the trained network. Approximately 93,000 type-1 particles were re-classified into the type-1 and type-1b classes. Each of the type-1 and type-1b particles were refined with an overall mask, and then subtracted with a mask around Rat1-Rai1 and the RNAPII stalk. 3D classifications of the subtracted particles were performed to exclude particles with poor Rat1-Rai1 density. 35,181 (type-1) and 41,265 (type-1b) particles with clear Rat1-Rai1 density were selected for the final reconstruction of the maps at 3.4 Å resolution. For the type-2 complex, approximately 140,000 particles were classified into apo RNAPII and RNAPII-Rat1-Rai1 complexes with different RNAPII clamp orientations. The class with the most (∼53,000) particles and the best map quality was selected for further refinement. The particles were initially refined with an overall mask, and then subtracted with a mask around Rat1-Rai1. 3D classifications of the subtracted particles were performed to exclude particles with poor Rat1-Rai1 density, and 49,931 particles with clear Rat1-Rai1 density were selected for the final reconstruction of a 3.4 Å resolution map.

### Model Building and Refinement

In the Rat1-Rai1-Rtt103 complexes, RNA was not observed, and in the RNAPII-Rat1-Rai1 complexes, Rtt103 was not observed. To build the five structures, the structures of *kp*Rat1(2Q3N), *kp*Rai1(2A), and *kp*Rtt103, which were generated by Alphafold2 ^54^, and *kp*RNAPII (PDB: 5XOG) were placed and rigid-body fitted into the cryo-EM map using UCSF Chimera ^55^. The models were manually built in WinCoot 0.9.8.7 ^56^ and ISOLDE ^57^ with the guidance of the cryo-EM map, and overall real-space refinement was performed using Phenix 1.9 ^58^. The data validation statistics are shown in Supplementary Table 1. The final model was validated with Molprobity ^59^ and Procheck ^60^. Figures were prepared using UCSF Chimera 1.17, ChimeraX, and PyMOL2.4.0 [http://pymol.sourceforge.net/].

### Reporting summary

Further information on research design is available in the Nature Portfolio Reporting Summary linked to this article.

## Data availability

The cryo-EM maps and coordinates have been deposited in the Electron Microscopy Data Bank (EMDB) and Protein Data Bank (PDB), respectively, with accession codes: EMD-39221 and 8YFE (Rat1-Rai1), EMD-39211 and 8YF5 (Rat1-Rai1-Rtt103N), EMD-39226 and 8YFQ (EC-Rat1-Rai1 (type 1)), and EMD-39227 and 8YFR (RNAPII-Rat1-Rai1 (type 2)).

## Acknowledgments

The cryo-EM experiments were performed with the Krios G4 microscope at the RIKEN Yokohama cryo-EM facility. This work was supported by Japan Society for the Promotion of Science KAKENHI Grant Number JP20H05690 (to S.S.).

## Author contributions

T.Y. and S.S. designed this study. T.Y., Y.M. and H.E. performed cryo-EM analyses. T.Y. performed protein preparations and biochemical analyses. M.G. prepared *K. phaffii* RNAPII. M.A. performed DNA cloning and mutagenesis. T.Y., Y.M., H.E. and S.S. wrote the manuscript. All of the authors discussed the results and commented on the manuscript.

## Competing interests

Authors declare that they have no competing interests.

## Notes

### Competing Interest Statement

The authors have declared no competing interest.

