## Supplementary materials for "Structural basis of eukaryotic transcription termination by the Rat1 exonuclease complex"

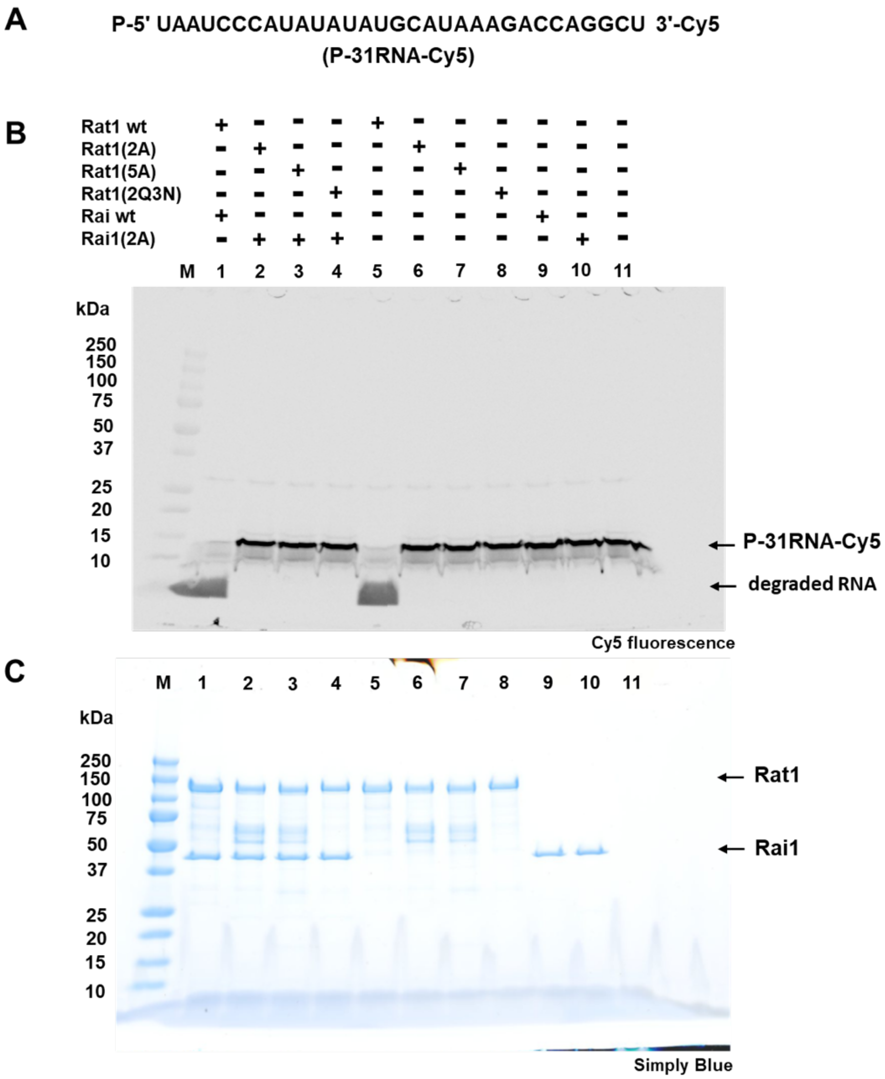


**Supplementary Figure 1.** ***In vitro*** **exonuclease activity of Rat1 and Rai1 from** ***K. phaffii***
Cleavage of 5´-monophosphorylated/3´-Cy5-labelled RNA substrate (P-31RNA-Cy5, A) by *kp*Rat1, *kp*Rai1, and *kp*Rat1-Rai1 complex. The Rat1-Rai1 exonuclease activities were measured *in vitro* as described in Materials and Methods. After reaction with wild-type Rat1 protein, more than 90% of P-31RNA-Cy5 was degraded, whereas no degradation of the RNA substrate was observed when reacting with mutant *kp*Rat1 proteins. Abbreviations are as follows. Rat1 wt: wild-type *kp*Rat1; Rat1(2A): *kp*Rat1(D233A/D235A); Rat1(5A): *kp*Rat1(E203A/E205A/D233A/D235A/D330A); Rat1(2Q3N): *kp*Rat1(E203Q/E205Q/D233N/D235N/D330N); Rai1(2A): *kp*Rai1(E213A/D215A). Reaction products were analyzed by SDS-PAGE, and visualized with Cy5 fluorescence (B), and Simply Blue-staining (C).

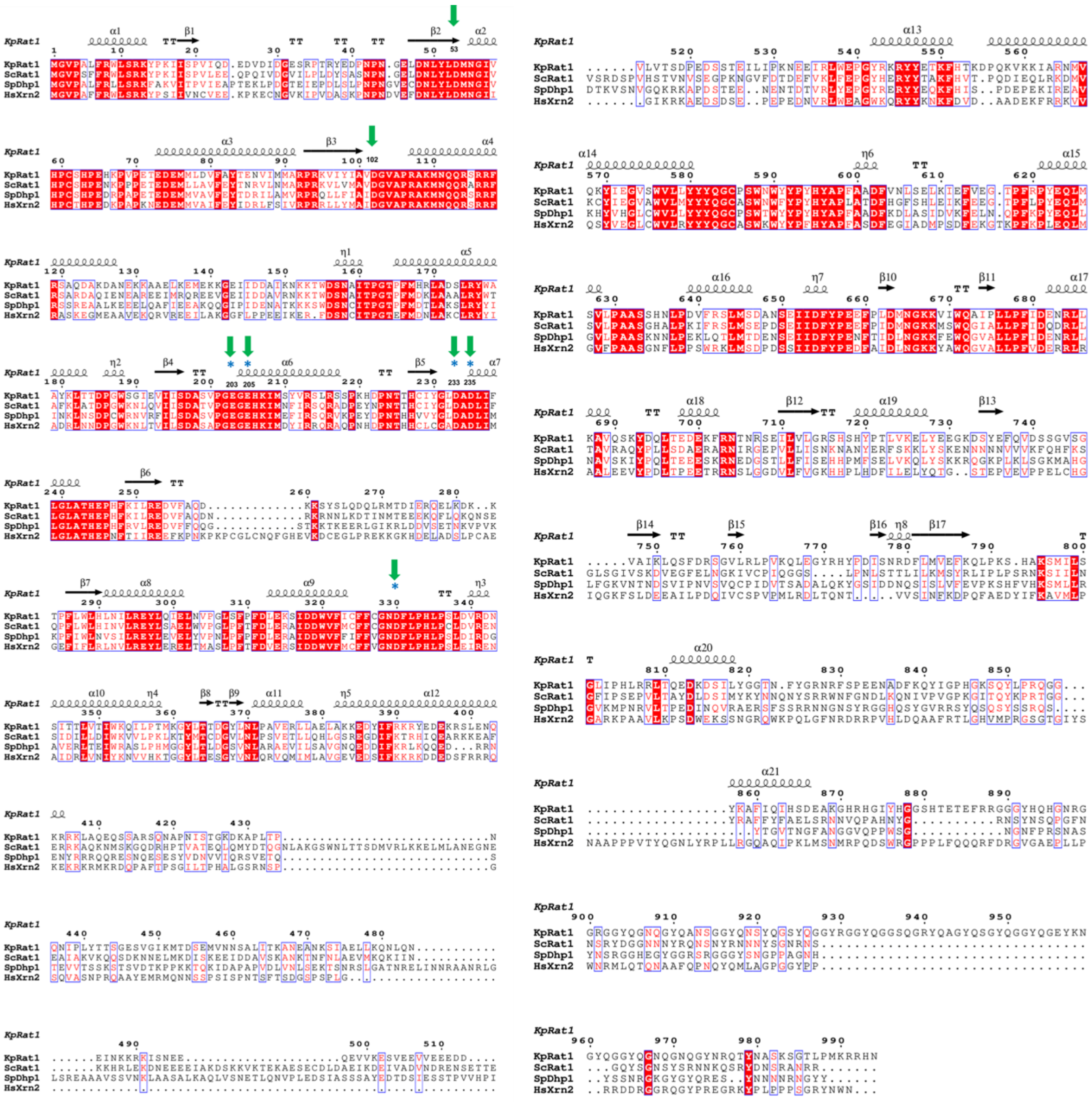

**Supplementary Figure 2. Sequence alignment of *kp*Rat1 and its orthologs.**

Sequence alignment of *K. phaffii* Rat1 (*Kp*Rat1, CCA39307) with orthologs from *S. cerevisiae* (*Sc*Rat1, CAA99240), *S. pombe* (*Sp*Dhp1, CAA93235), *H. sapiens* (*Hs*Xrn2, NP_036387). Amino acid sequences of Rat1 were aligned with CLUSTAL W [61], and the figure was prepared with ESPript [62]. Sequence identities of *kp*Rat1 with *sc*Rat1, *sp*Dhp1, and *hs*Xrn2, are 50, 46, and 43%, respectively. The conserved acidic residues (Asp53, Asp102, Glu203, Glu205, Asp233, Asp235, and Asp330) in the active site are highlighted with green arrows above the sequence alignment. Mutated amino acid residues (Glu203, Glu205, Asp233, Asp235, and Asp330) in the present study are highlighted as cyan asterisks.

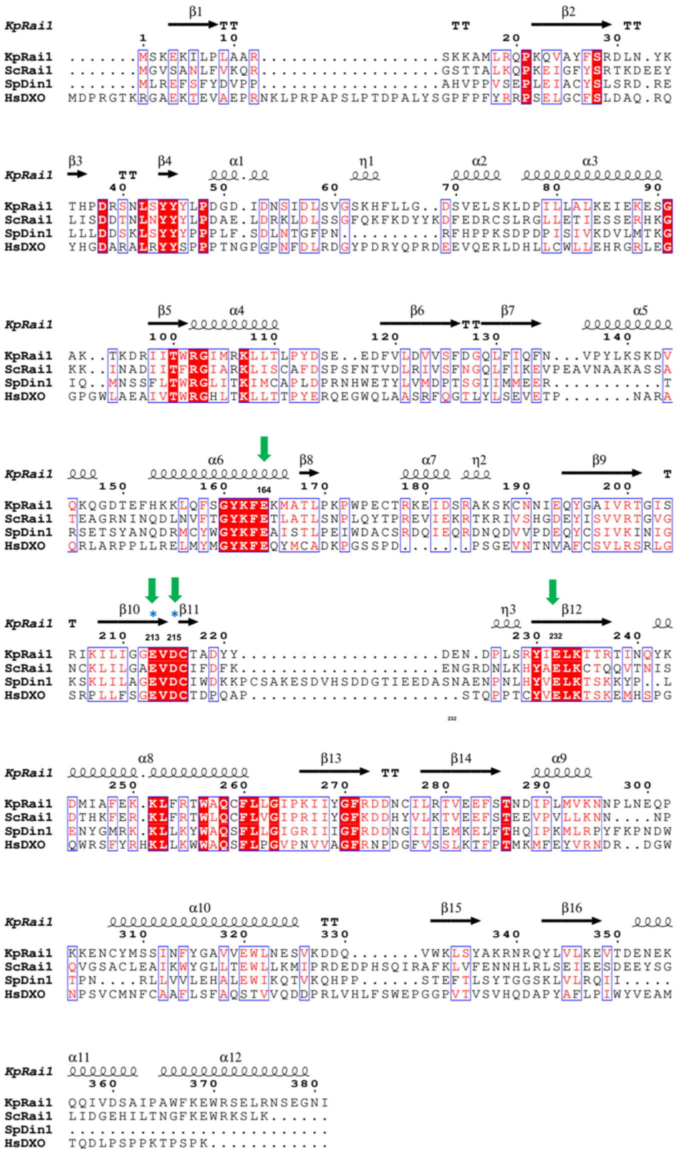

**Supplementary Figure 3. Sequence alignment of *kp*Rai1 and its orthologs.**

Sequence alignment of *K. phaffii* Rai1 (*Kp*Rai1, CCA37187) with orthologs from *S. cerevisiae* (*Sc*Ra11, KAF4004777), *S. pombe* (*Sp*Rai11, CAB16716), *H. sapiens* (*Hs*DXO, AQY76800). Amino acid sequences were aligned with CLUSTAL W, and the figure was prepared with ESPript. Sequence identities of *kp*Rai1 with *sc*Rai1, *sp*Rai1, and *hs*DXO, are 37, 28, and 27%, respectively. The conserved acidic residues (Glu164, Glu213, Asp215, and Glu232) in the active site are highlighted with green arrows above the sequence alignment. Mutated amino acid residues (Glu213, and Asp215) in the present study are highlighted as cyan asterisks.

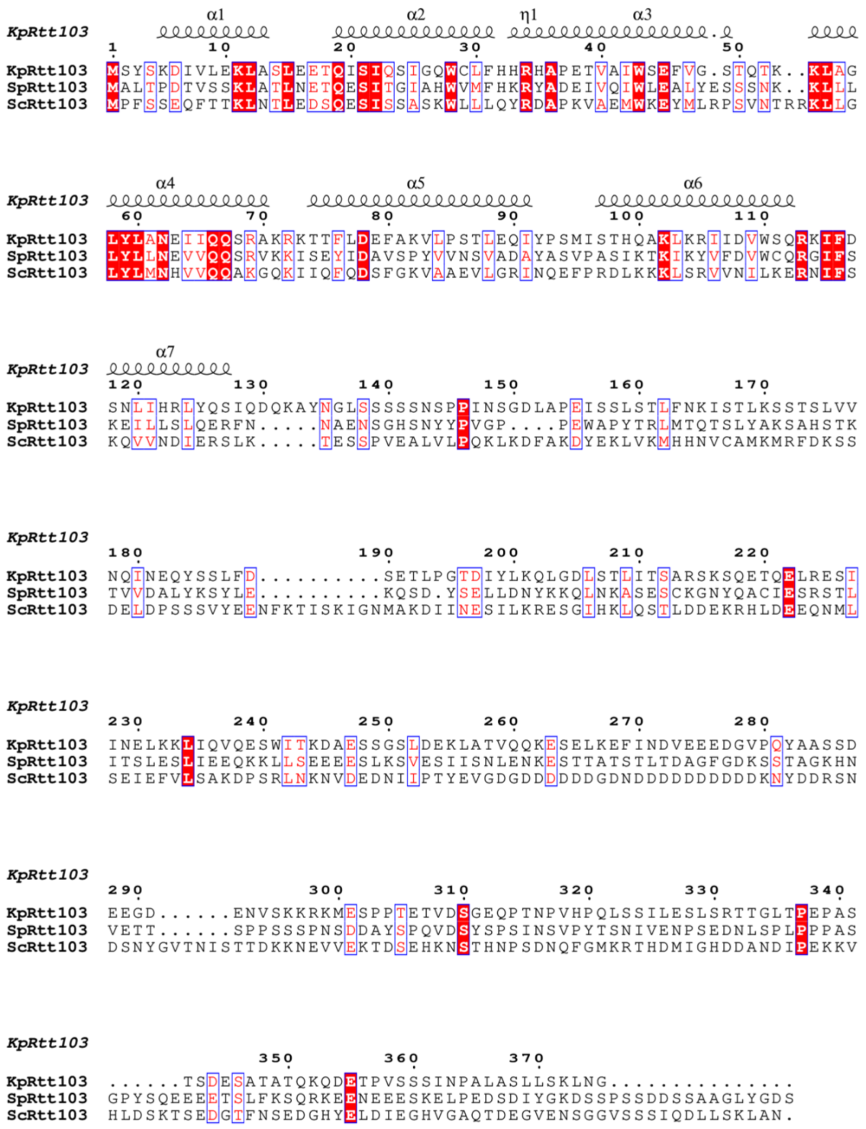


**Supplementary Figure 4. Sequence alignment of *kp*Rtt103 and its orthologs.**

Sequence alignment of *K. phaffii* Rtt103 (*Kp*Rtt103, CCA37063) with orthologs from *S. cerevisiae* (*Sc*Rtt103, NP_010575), *S. pombe* (*Sp*Rtt103, NP_595404). Amino acid sequences were aligned with CLUSTAL W, and the figure was prepared with ESPript. Sequence identities of *kp*Rtt103 with *sc*Rtt103, *sp*Rtt103 are 20% and 32%, respectively. There are no Rtt103 orthologs in *H. sapiens*.

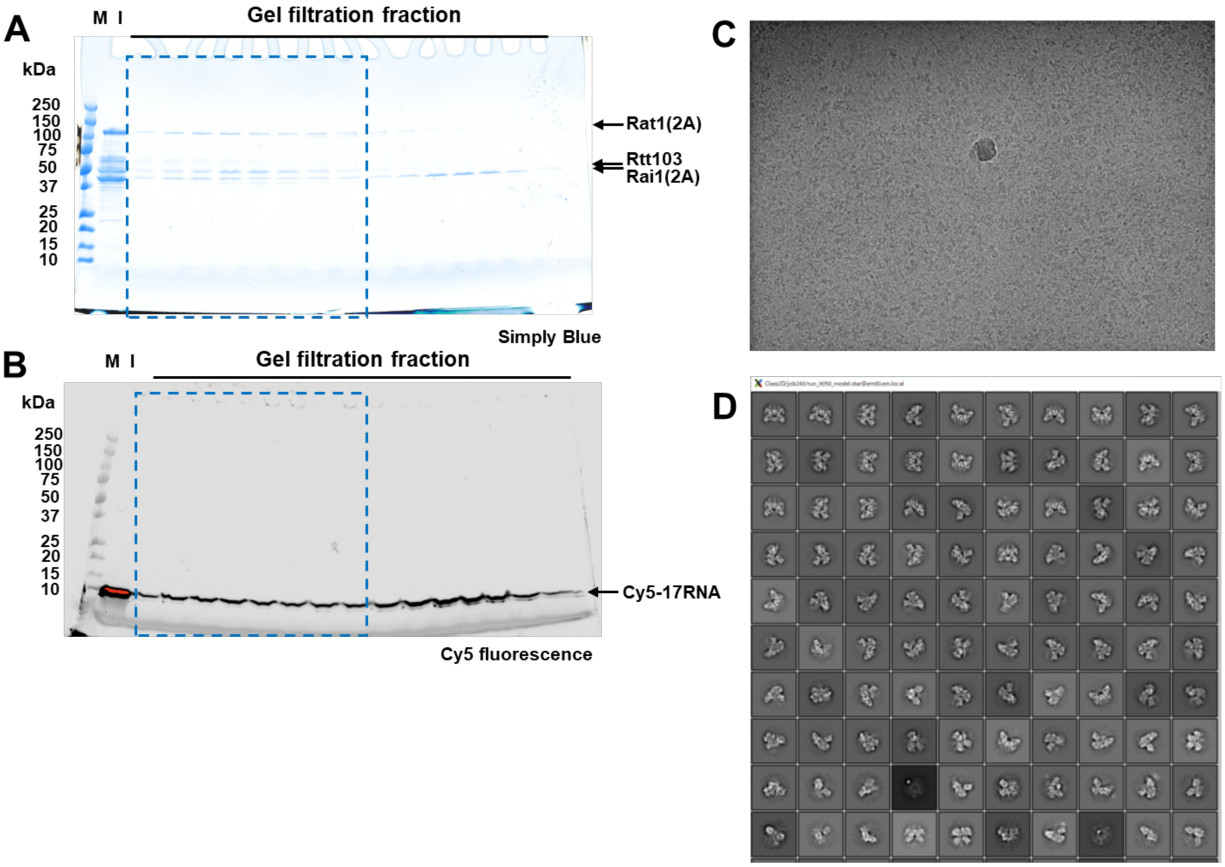


**Supplementary Figure 5. Sample preparation and image analyses of the Rat1-Rai1-Rtt103 complex.**

(A, B) Electrophoretic analyses of gel filtration fractions. Proteins and Cy5-RNA were detected by Simply Blue-staining (A) and Cy5 fluorescence (B), respectively. Fractions marked with blue boxes were collected for cryo-EM sample preparation. M: molecular mass standards; I: loaded protein samples for the gel-filtration experiments. (C) A raw micrograph of a cryo-EM grid used in the present study. (D) 2D class averages for the Rat1-Rai1 tetramer complex after particle picking by TOPAZ.

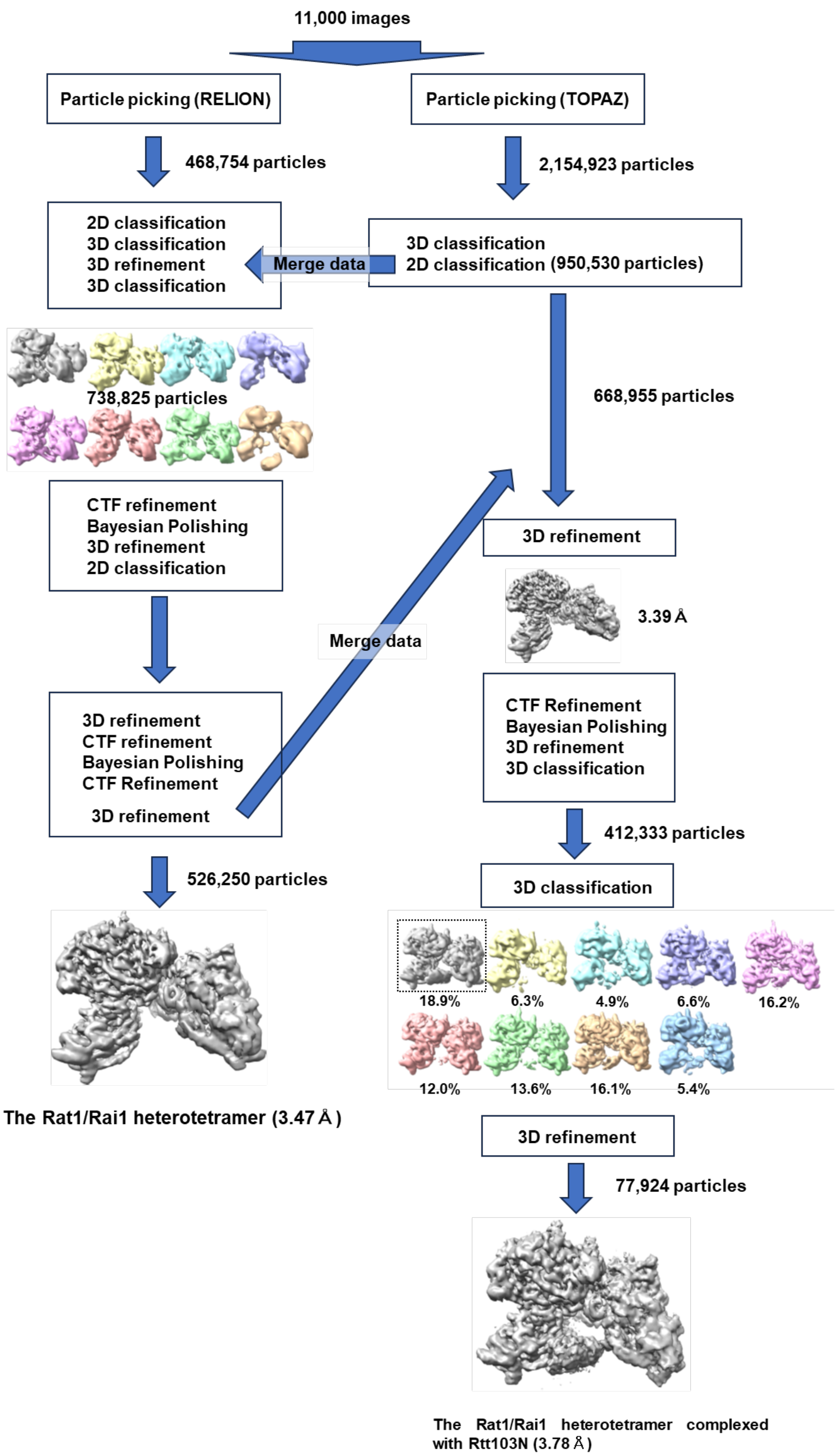


**Supplementary Figure 6**. **Cryo-EM data processing workflow for the Rat1-Rai1-Rtt103 complex.**

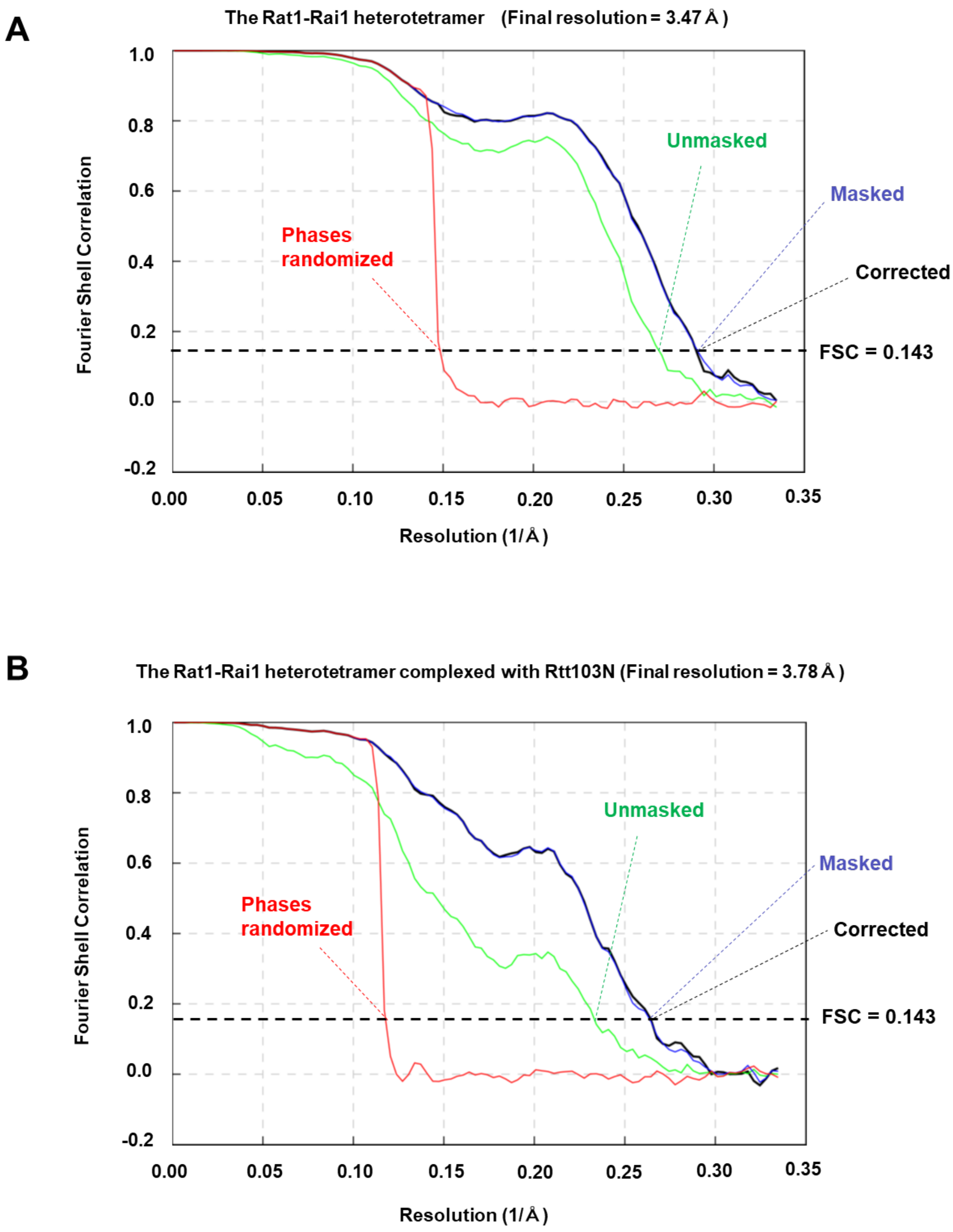


**Supplementary Figure 7. FSC curves of the Rat1-Rai1 heterotetramer complex (A) and the Rat1-Rai1-Rtt103N heteropentamer complex (B).**

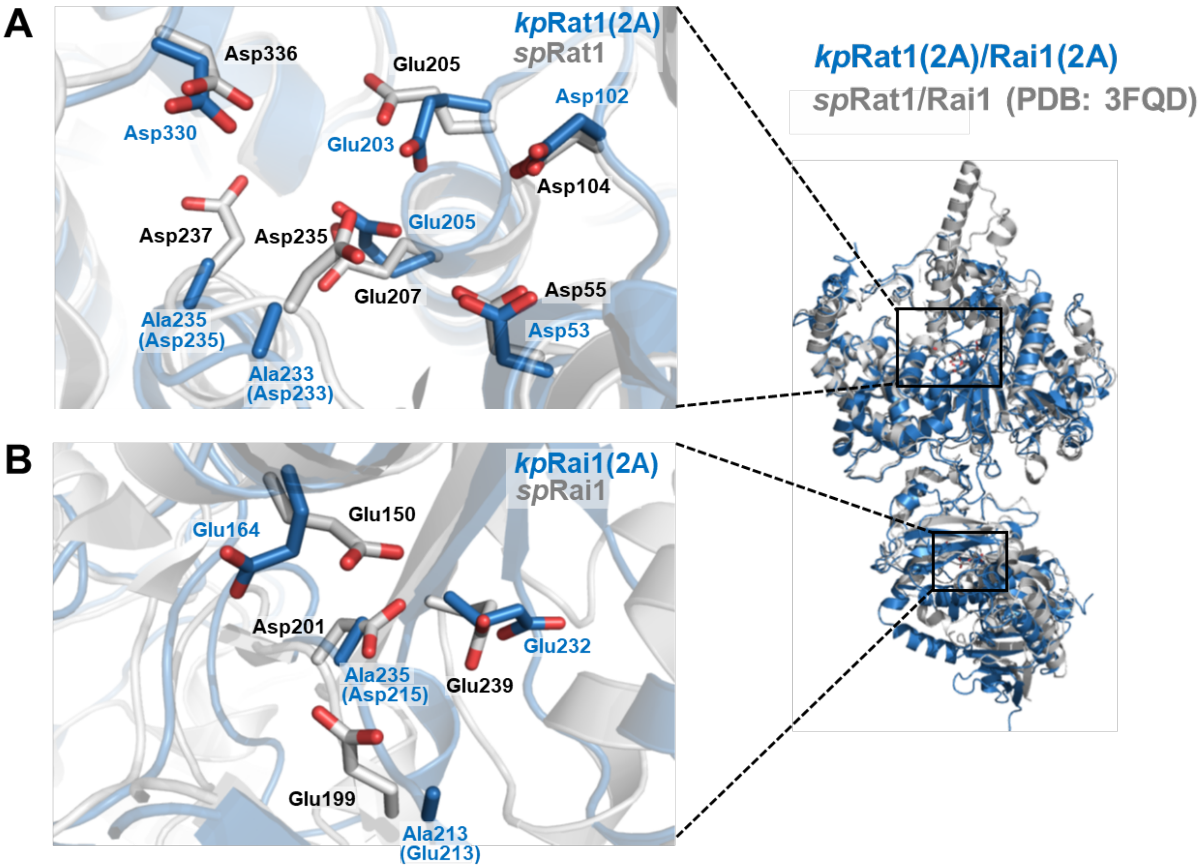


**Supplementary Figure 8. Active-site structures of Rat1 and Rai1 from *K. phaffii*.**
Close-up views and comparison of the Rat1 and Rai1 active site residues (shown as stick models) with those of *sp*Rat1-Rai1. The active site residues of *kp*Rat1 and *kp*Rai1 (blue) were superimposed well on the *sp*Rat1 and *sp*Rai1 structures (white). The conserved acidic residues of *kp*Rat1(2A) [Asp53, Asp102, Glu203, Glu205, Ala233(Asp233), Ala235(Asp235), and Asp330] and those of *sp*Rat1 (Asp55, Asp104, Glu205, Glu207, Asp235, Asp237, and Asp336) are shown as blue and white stick models, respectively (A). The conserved acidic residues of *kp*Rai1(2A) [Glu164, Ala213(Glu213), Ala215(Asp215), and Glu232] and those of *sp*Rai1 (Glu150, Glu199, Asp201, and Glu239) are shown in blue and white stick models, respectively. (B) Overall ribbon models of *kp*Rat1/Rai1 superimposed on that of *sp*Rat1-Rai1 (PDB: 3FQD) were shown in the right.

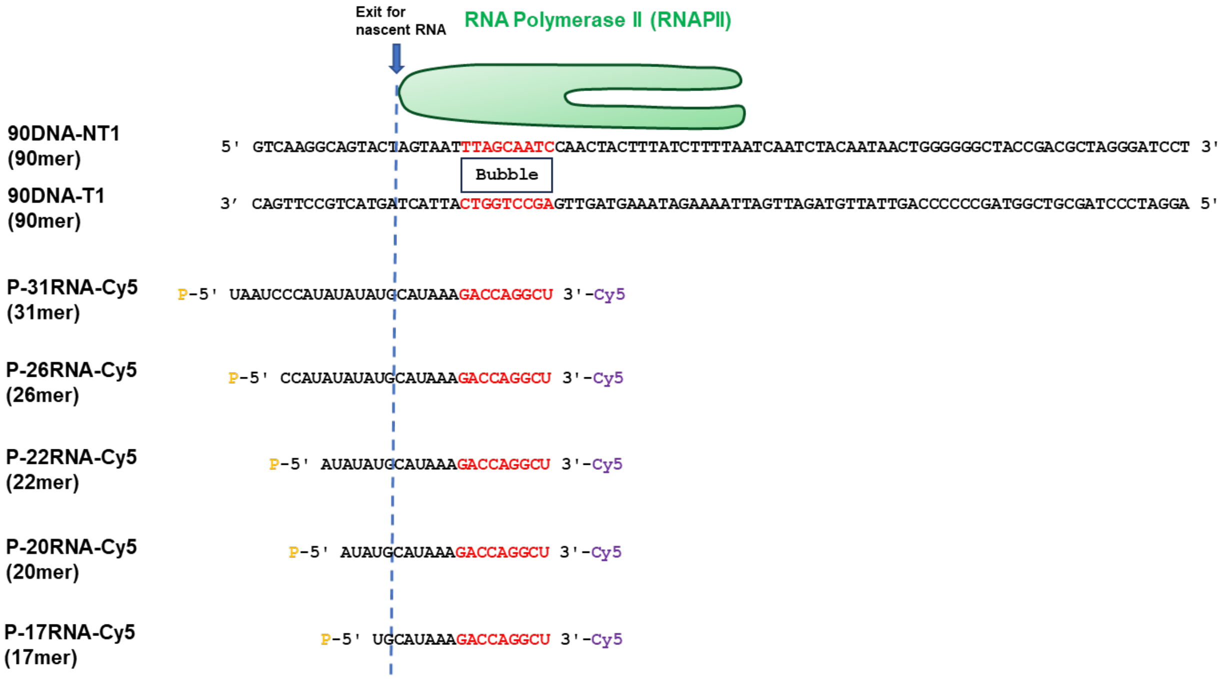


**Supplementary Figure 9. DNA and RNA sequences used for the RNAPII-Rat1-Rai1 complex assembly.**

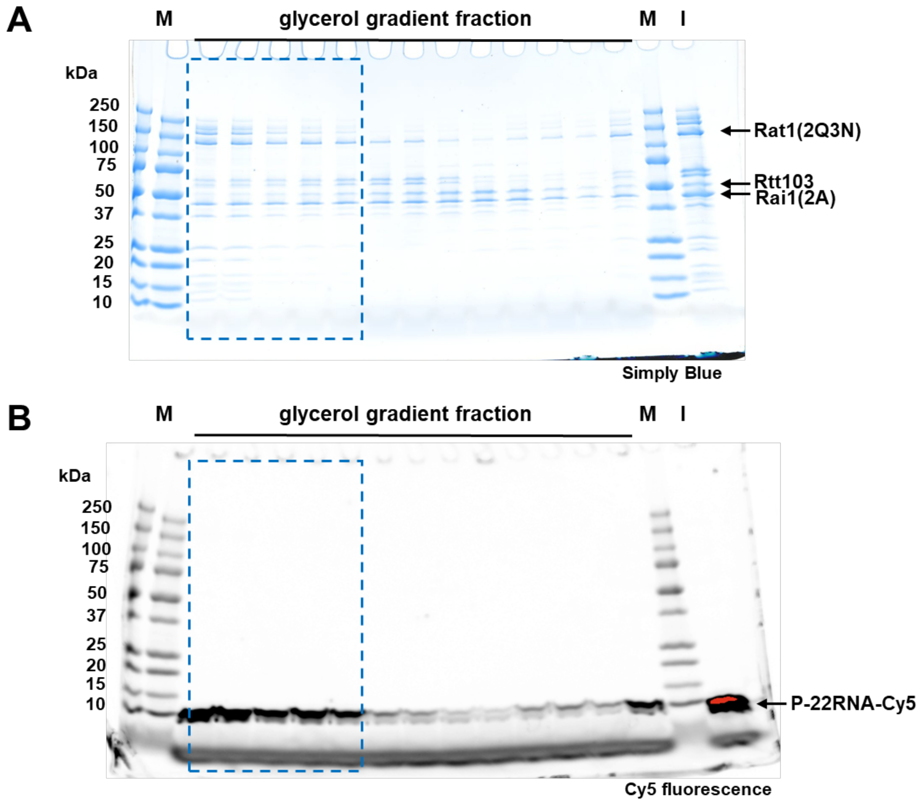


**Supplementary Figure 10. Sample preparation and image analyses of the RNAPII-Rat1-Rai1 complexes.**

(A, B) Electrophoretic analyses of glycerol gradient fractions. Proteins and P-22RNA-Cy5 were detected by Simply blue-staining (A), and Cy5 fluorescence (B), respectively. Fractions marked with blue boxes were collected for cryo-EM sample preparation. M: molecular mass standards; I: loaded protein samples for the glycerol gradient experiments.

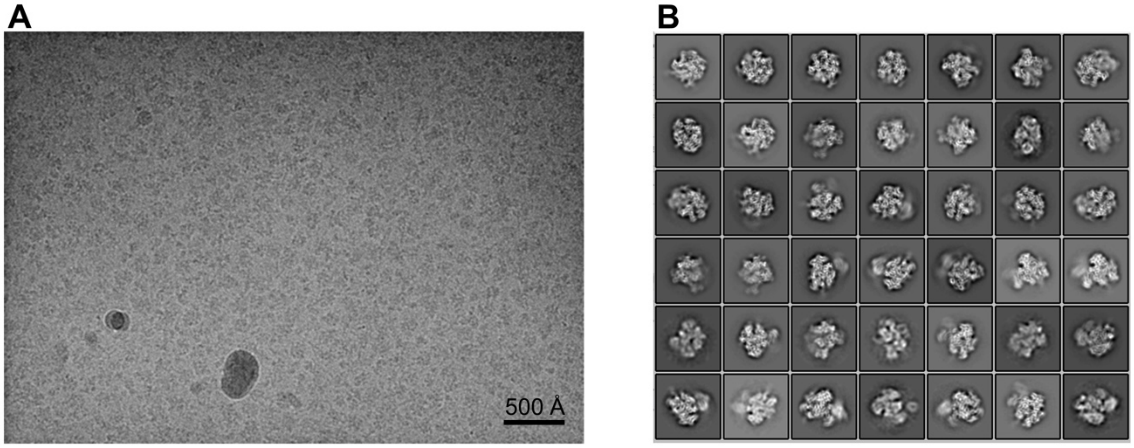


**Supplementary Figure 11. Image analysis of the RNAPII-Rat1-Rai1 complexes.**
(A) A raw micrograph of a cryo-EM grid. (B) 2D class averages for the RNAPII and RNAPII-Rat1-Rai1 complexes.

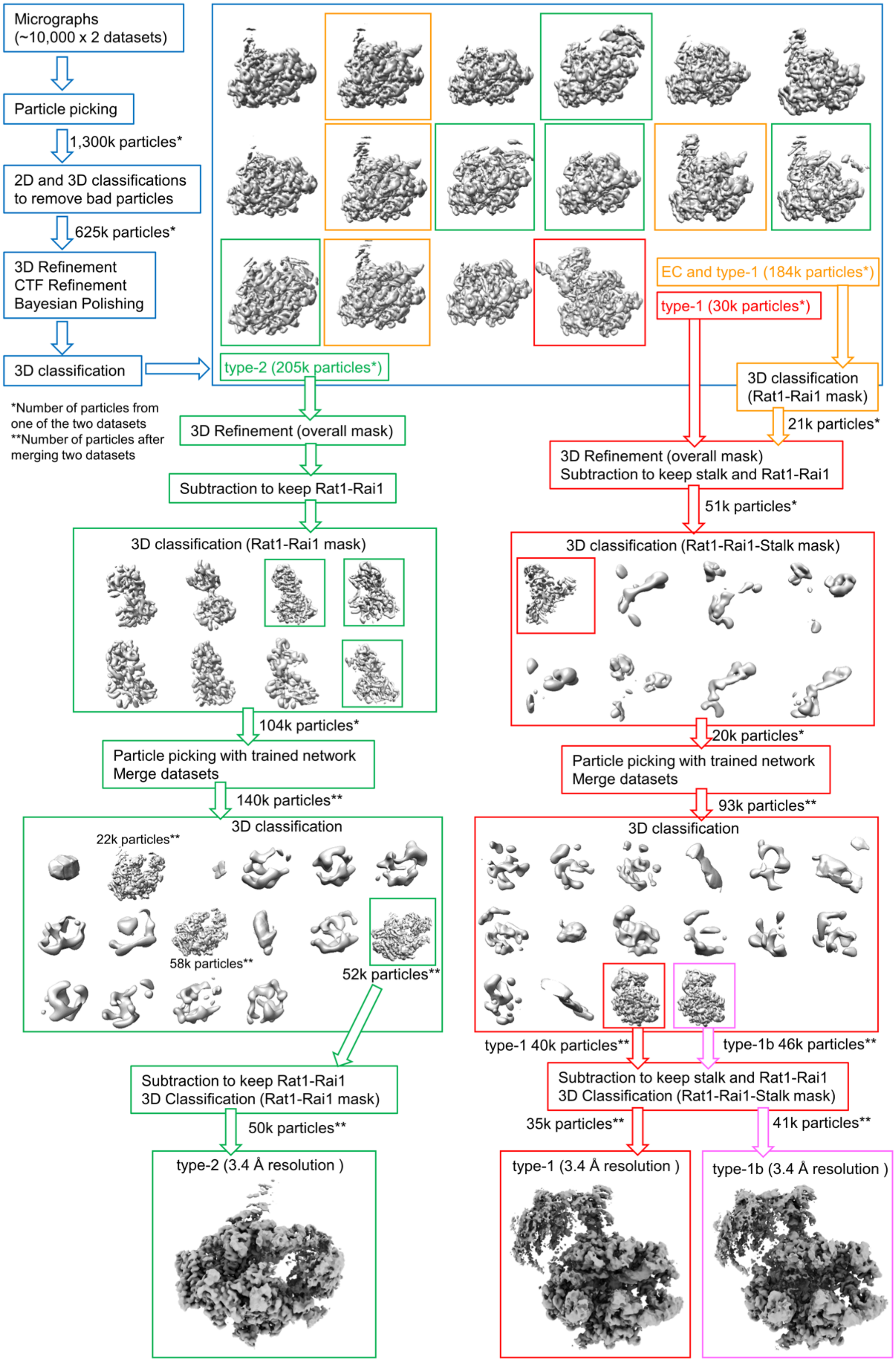


**Supplementary Figure 12. Cryo-EM data processing workflow for the RNAPII-Rat1-Rai1 complexes.**

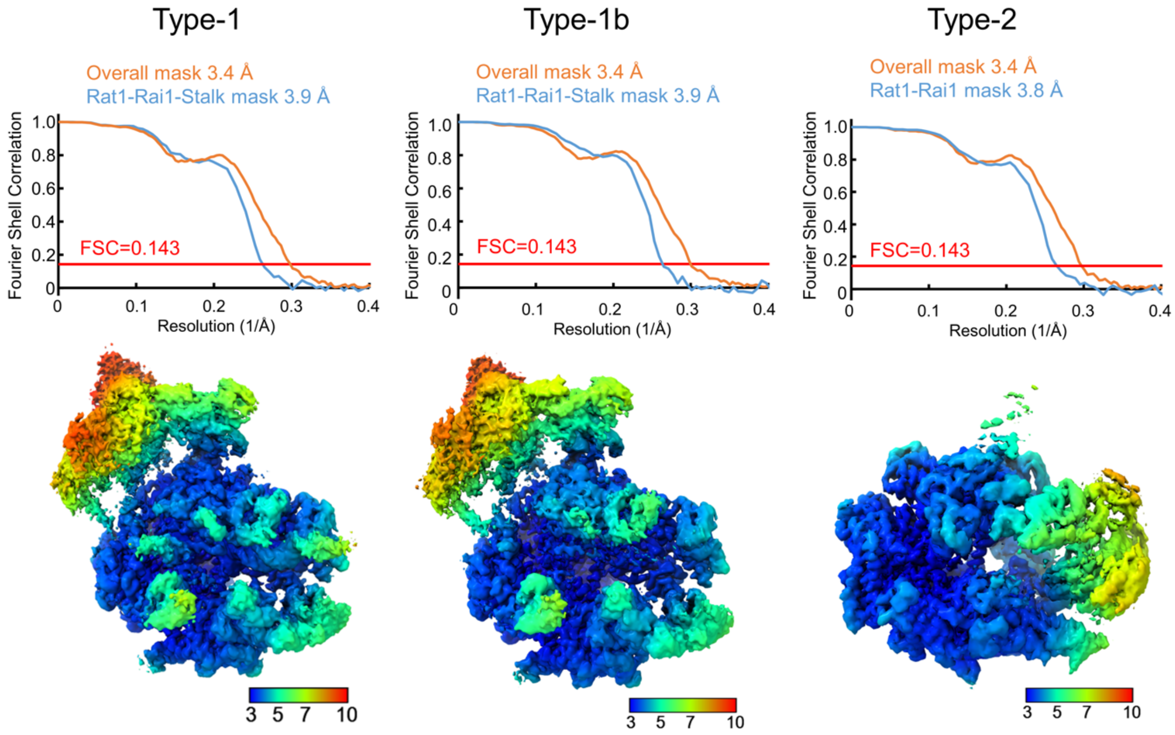


**Supplementary Figure 13. Gold-standard Fourier shell correlation (FSC) curves and local resolution maps of the RNAPII-Rat1-Rai1 complexes.**

Top, FSC curves for the type-1 (left), type-1b (middle) and type-2 (right) complexes. FSC curves calculated using overall masks and local masks around Rat-Rai1(-stalk) are shown in orange and blue, respectively. Bottom, cryo-EM maps colored by local resolutions estimated by Relion.

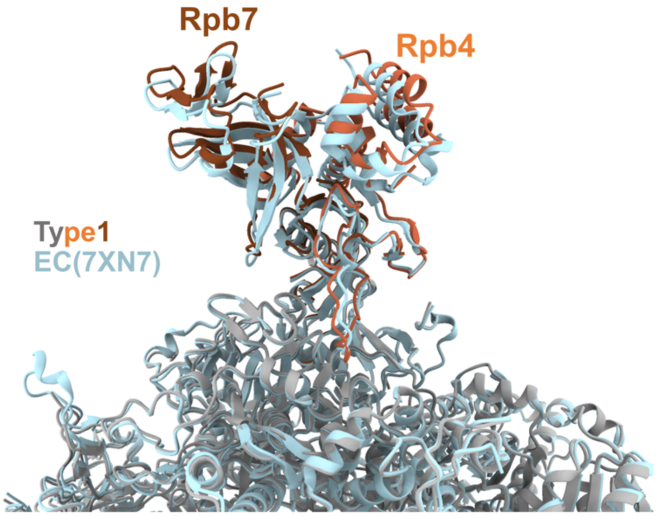


**Supplementary Figure 14. Comparison of the RNAPII stalk conformation between the type-1 complex and EC.**

Structure of RNAPII EC (PDB 7XN7; light blue) is superimposed with the type-1 complex (gray) by the Rpb1 subunit. The Rpb4 and Rpb7 subunits of the type-1 complex are colored orange and brown, respectively.

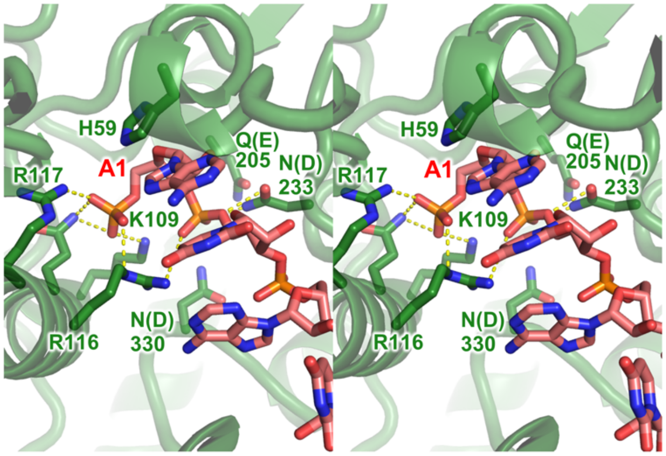


**Supplementary Figure 15. Structure of the Rat1 active site incorporating the RNA 5′-end.**

The Rat1 active site structure in the type-1 complex is shown in a wall-eye stereo view.

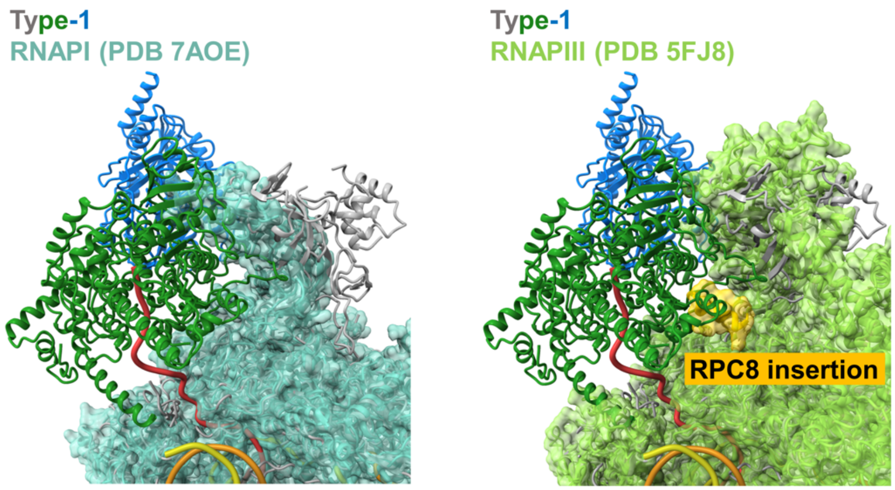


**Supplementary Figure 16. Superimposition of the type-1 complex with RNAPI and RNAPIII EC structures**

Structures of RNAPI EC (PDB 7AOE; cyan) and RNAPIII EC (PDB 5FJ8; yellow green) are superimposed on the type-1 complex (colored as in Fig. 2) by the Rpb1 subunit. The specific insertion in the RPC8 (corresponding to Rpb7 in RNAPII) is colored yellow.

### Supplementary Table 1. Data collection and refinement statistics

| **Data collection** | **Rat1-Rai1-Rtt103** | | **RNAPII-Rat1-Rai1 dataset 1** | **RNAPII-Rat1-Rai1 dataset 2** |
|---|---|---|---|---|
| EM equipment | Krios G4 | | Krios G4 | Krios G4 |
| Magnification | 105,000 | | 105,000 | 105,000 |
| Voltage (kV) | 300 | | 300 | 300 |
| Detector | Gatan K3 | | Gatan K3 | Gatan K3 |
| Pixel size (Å) | 0.83 | | 0.83 | 0.83 |
| Electron dose ($e^-$/Å$^2$) | 61.9 | | 53.3 | 53.3 |
| Defocus range (µm) | –1.6 ~ –2.4 | | –1.6 ~ –2.4 | –1.6 ~ –2.4 |
| Number of micrographs | 11,000 | | 10,001 | 11,134 |
| | **Rat1-Rai1** | **Rat1-Rai1-Rtt103** | **EC-Rat1-Rai1 (type-1)** | **RNAPII-Rat1-Rai1 (type-2)** |
| **Refinement** | | | | |
| Resolution (Å) | 3.5 | 3.8 | 3.4 | 3.4 |
| Final number of particles | 526,250 | 77,924 | 35,181 | 49,931 |
| Symmetry Imposed | C2 | C1 | C1 | C1 |
| RMS(bonds) (Å) | 0.002 | 0.003 | 0.003 | 0.002 |
| RMS(angles) (°) | 0.537 | 0.524 | 0.557 | 0.545 |
| Ramachandran plot<br>favored (%)<br>allowed (%)<br>outliers (%) | <br>95.62<br>4.38<br>0.00 | <br>95.36<br>4.71<br>0.00 | <br>96.61<br>3.31<br>0.08 | <br>96.11<br>3.77<br>0.12 |
| Validation<br>Molprobity score<br>Clashscore<br>Poor rotamers (%) | <br>1.85<br>10.71<br>0.05 | <br>1.88<br>10.84<br>0 | <br>1.64<br>7.78<br>0.18 | <br>1.78<br>9.81<br>0 |
| Number of non-hydrogen atoms | 17,315 | 18,476 | 42,465 | 39,461 |
| Protein residues | 2,107 | 2,250 | 5,070 | 4,921 |
| DNA/RNA residues | 0 | 0 | 88 | 0 |
| ligands | 0 | 0 | 9 | 9 |
| PDB ID | 8YFE | 8YF5 | 8YFQ | 8YFR |
| EMDB ID | 39221 | 39211 | 39226 | 39227 |
